# Large-scale Dynamical Fingerprints of Distributed Synaptic Alterations in Early-Stage Psychosis

**DOI:** 10.1101/2024.07.15.603513

**Authors:** Ayelet Arazi, Alessandro Toso, Tineke Grent-‘t-Jong, Peter J. Uhlhaas, Tobias H. Donner

## Abstract

Psychotic disorders, such as schizophrenia, present a major challenge for research and clinical practice: its pathogenesis is complex, the individual symptomatology is heterogenous, and there is a lack of biomarkers for the early detection, diagnosis, and individualized treatment. Mounting evidence indicates that the synaptic and microcircuitry alterations underlying psychosis are widely distributed in the brain. Here, we developed a magnetoencephalography approach to map the resulting alterations of local cortical population dynamics across the human cerebral cortex. We identified large-scale patterns of changes in neural dynamics that were remarkably similar between first-episode psychosis patients and individuals at clinical high risk for psychosis. These spatial patterns also resembled those induced by pharmacological manipulations of excitatory NMDA glutamate receptors and inhibitory GABA-A receptors in healthy participants. Differences in those spatial patterns of cortical dynamics between psychosis patients related to individual symptomatology. Our study opens a new window on the distributed pathophysiology of psychosis.

**Teaser:** Cortex-wide changes in neural mass dynamics due to psychosis resemble changes due to GABA-A or NMDA receptor manipulation and relate to clinical symptoms.

## Introduction

Psychotic disorders, such as schizophrenia, present a major burden to society (*1–3*). Schizophrenia is characterized by positive and negative symptoms as well as by cognitive deficits, which may originate from diverse underlying mechanisms (4). Aberrations in several neurotransmitter systems have been implicated in schizophrenia, including disturbances of GABAergic cortical inhibition (5–8) and glutamatergic excitation, specifically via NMDA receptors (9, 10). Schizophrenia is considered a neuro-developmental disorder, with aberrations in brain circuitry emerging long before symptoms typically manifest during adolescence (3). Recently, criteria have been developed to allow the detection of individuals at clinical high-risk for psychosis (CHR-P) (11). Approximately 25% of CHR-P individuals transition to psychosis within 3 years of follow-up (12). Accordingly, one important objective of current research is to identify biomarkers for psychotic disorders, as well as for prediction and prognosis in CHR-P participants (13–17).

Cortical circuits exhibit an intricate arrangement of excitatory and inhibitory synaptic connections. The detailed properties of cortical circuits, such as the ratio between excitatory and inhibitory connections, shape the mechanisms underlying cognitive behaviors that involve working memory, accumulation of evidence, and deliberative decision-making (18–26). These microcircuit properties are disturbed in schizophrenia and related disorders, such as autism, whereby a major focus has been on the local ratios between synaptic excitation and inhibition within cortical areas (9, 10, 18, 27–30). Simulations of neural circuit models show that changes in such microcircuit properties produce characteristic changes in the dynamics of spontaneous neural mass action, as expressed in local field potentials (31–37). Because such neural mass signals are detectable with non-invasive electrophysiological recording techniques (such EEG or MEG) in humans, this insight holds promise for the development of biomarkers for psychotic disorders (29, 38).

Recent advances have shown that properties of cortical microstructure, including synaptic inhibition via GABA-A receptors and excitation via NMDA receptors, are not homogeneous but exhibit gradients across cortex, which reflect the anatomical cortical hierarchy (22, 39). Similar gradients exist for features of cortical dynamics (22, 34, 40–43). Mounting evidence indicates that the circuit dysfunctions underlying psychotic disorders (3, 29) are also expressed across many cortical areas. How such distributed alterations in (microscopic) circuit properties translate into changes in large-scale cortical dynamics and into psychotic symptoms is currently unknown. We hypothesized that there may be characteristic large-scale patterns of altered cortical dynamics in psychosis, which result from alterations in GABA-A or NMDA receptor function. To test this hypothesis, we developed a non-invasive MEG approach to (i) identify putative large-scale neurophysiological signatures of early-stage psychosis and (ii) unravel their underlying synaptic mechanisms.

We comprehensively mapped the cortex-wide patterns of psychosis-related alterations of spontaneous cortical population dynamics (across 180 well-defined areas per hemisphere). Inspired by recent advances in large-scale computational neuroanatomy (39, 43–45) and clinical neurophysiology (46, 47), we developed a combination of MEG source-imaging of cortical population dynamics (48), theory-driven selection of a set of dynamics parameters (see Results for details) (41, 49), and multivariate pattern analyses. This approach uncovered highly similar large-scale patterns of changes in cortical dynamics in first-episode psychosis patients and CHR-P individuals. These patterns, in turn, resembled the patterns of changes in cortical dynamics induced by manipulating GABA-A or NMDA receptor function in healthy participants. Differences in those spatial patterns of cortical dynamics between first-episode psychosis patients related to individual symptomatology.

## Results

We analyzed resting-state MEG data collected in two previously published studies (13–16, 50, 51). The clinical dataset was collected at the University of Glasgow in a group of CHR-P participants (N=117), a group of first episode psychosis patients (FEP; N=32), and a group of healthy controls (HC; N=45; see Methods for details on the samples). The pharmacological dataset was collected at the University Medical Center Hamburg-Eppendorf and measured the effects of double-blind oral administration of the GABA-A agonist lorazepam (LZP) and the NMDA receptor antagonist memantine (MEM), each compared to placebo within a group of 20 healthy participants (cross-over design, see Methods). All groups completed a block of eyes-open resting-state, fixating a central cross on an otherwise blank screen.

### A six-parameter assay of local cortical population dynamics

We parameterized spontaneous cortical network dynamics in the clinical and pharmacological datasets in terms of a set of six parameters (Figure 1 and Methods). The power spectra of local MEG signal fluctuations were decomposed into oscillatory (periodic) and non-oscillatory (aperiodic) components (Figure 1A left; Methods and (52)), from which we extracted a total of five parameters (37): the (i) exponent, (ii) “knee frequency”, and (iii) area under-the-curve (of aperiodic fit only) of the aperiodic component. We further extracted the (iv) power and (v) peak frequency specifically from the periodic fit to the alpha (7-12 Hz) frequency range. In a separate analysis (Figure 1A right; Methods), we quantified the so-called (vi) Hurst exponent, quantifying long-range temporal correlations in the fluctuations of MEG signal amplitude envelopes, also focusing on in the alpha band (53).

**Figure 1:**
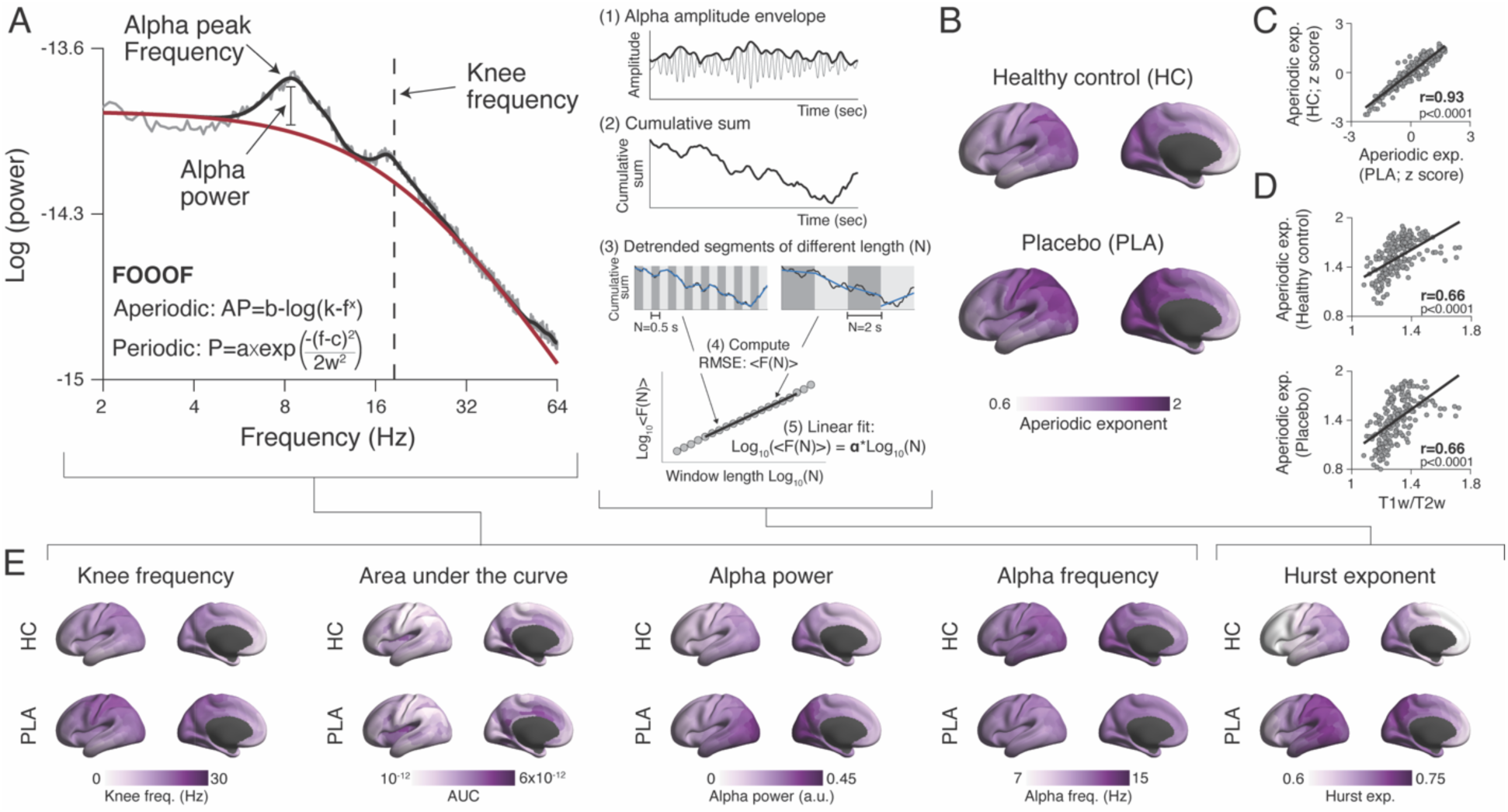
Large-scale spatial patterns of local cortical population dynamics. **(A)** Decomposing power spectra of cortical population activity. Left: Power spectral density (PSD, gray) of resting-state MEG signal in an example data set (single-participant, V1; placebo condition). Black and red lines are the fitted periodic and aperiodic components, respectively (Methods). Right: Illustration of detrended fluctuation analysis of alpha-envelope fluctuations: (1) Alpha (8-13Hz) amplitude envelope of MEG signal. (2) Cumulative sum of amplitude envelope. (3) Detrended segments of cumulative sum, shown for two window lengths N (0.5s or 2s). (4) Fluctuation function <F> as a function of window lengths N, in log-log coordinates. (5) Scaling (Hurst) exponent, α, as the slope of the linear fit. **(B)** Cortical spatial distribution of the aperiodic exponent across 180 areas (left and right hemispheres averaged). (**C)** Across-area correlations between the maps of aperiodic exponent in both samples shown in panel B. Spearman’s correlation coefficients and p-values (permutation tests; corrected to account for spatial autocorrelations) are noted in each panel. Each dot represents a single parcel. Black line: linear fit. **(D)** Spatial similarities (across-area correlations) between the maps of aperiodic exponents (for each dataset) and of a structural MRI maker of cortical hierarchy (the ratio between T1w to T2w images). **(E)** Cortical spatial distribution of five additional parameters of cortical dynamics. Top row, healthy control (HC). Bottom row, placebo condition (PLA).

The selection of the above parameters was theory-driven, because these parameters have been related to key microcircuit properties, which may be altered in psychosis (3, 6, 9, 10, 28, 29) and other mental disorders (25, 27, 54, 55). Specifically, the aperiodic exponent of the overall power spectrum and Hurst exponent of alpha-envelope fluctuations have been related to the ratio between local synaptic excitation and inhibition (32, 33, 35–37). The knee frequency has been linked to the intrinsic timescale of cortical activity (34, 56), which, in turn, depends on the strength of recurrent excitation (22). The area under the curve of the aperiodic fit is a proxy of population firing rates (31). Work on neural oscillations has specifically implicated gamma-band responses as a marker of excitatory-inhibitory interactions within cortical microcircuits (*36*, *57–59*) and of aberrations thereof in psychosis (*14*, *29*). For quantification of periodic cortical activity (i.e., parameters (iv) to (vi)), we here focused on the alpha-range rather than gamma-range range. This was because our approach relied on large-scale (across-area) spatial patterns and alpha peaks are generally more prominent and widely distributed at rest (*58*), and were more consistently detected in our data across cortical areas and participants. Please note that the aperiodic parameters (i) to (iii) were based on the entire frequency range of the power spectra analyzed here, so our parameter set provided a rich summary of cortical dynamics.

### Large-scale spatial patterns of local cortical dynamics

The above six parameters of local cortical dynamics (henceforth called “dynamics parameters”) were comprehensively mapped across 180 well-defined areas covering the cortical surface (*44*). Throughout, we averaged across corresponding areas from the left and right hemispheres and visualized the resulting parameter maps on the surface of the left hemisphere. This approach enabled us to characterize large-scale spatial patterns of cortical dynamics in CHR-P and FEP-groups versus control groups, and under pharmacological interventions versus placebo conditions in healthy individuals.

All dynamics parameters exhibited a heterogenous spatial pattern, with a distinctive gradient across areas (Figure 1B; E). This gradient reflected the gradient of an MRI-based inverse proxy of myelination and anatomical hierarchy, the T1w/T2w ratio (*39*), across cortical areas (Figure 1D; Figure S1A, B). For example, the spatial patterns of the exponent of the aperiodic component, a putative measure of local excitation-inhibition ratio (*33*), were both positively correlated with the T1w/T2w ratios (Figure 1D: r(180)=0.66, p<10^-4^; spatial autocorrelation-preserving permutation tests; Methods), and they were also strongly correlated among one another (Figure 1C: r(180)= 0.93, p<10^-4^). In other words, the exponent of the aperiodic component became systematically and reliably smaller across the cortical hierarchy, with larger exponents for early sensory cortices and smaller exponents for prefrontal cortex. Likewise, the maps of the knee frequency, an inverse measure of intrinsic neural timescales (*34*, *56*) showed a decrease of knee frequencies from sensory to association cortex, in line with slower neural timescales in higher-tier areas (*22*, *40*, *41*).

The maps of the other parameters (Figure 1E) were, likewise, correlated with the T1w/T2w ratio maps for all but one (area-under-the-curve) in both datasets (Figure S1 A, B) and exhibited robust correlations among one another (Figure S1 D, E). This was the case in the data from both studies, whereby the spatial patterns of all parameters except of alpha peak frequency were strongly correlated between these two samples (Figure S1C). Importantly, similar spatial correlations to T1w/T2w ratios were evident in both clinical groups and both pharmacological conditions (Figure S1 F, G). Thus, the relationship between circuit dynamics and cortical hierarchy is overall robust to rapid (within-session) manipulation of important neurotransmitter receptors or longer-term, clinical aberrations of circuit function.

In sum, our results so far replicate the insight previously established with invasive recording techniques (*34*, *40*) that parameters of cortical dynamics (specifically, the knee frequency) reflect an important large-scale organizing principle of the brain, the cortical hierarchy. Our results go beyond previous findings in showing that the cortex-wide spatial patterns of cortical dynamics (i) are highly reproducible across independent samples obtained in separate MEG laboratories, and (ii) reflect cortical hierarchy also in patients with early-stage psychosis, indicating that the functional cortical hierarchy is preserved in this clinical condition.

### Dynamics parameters alterations in psychosis reflect GABA-A receptor densities

Having established the reliability and functional significance of the overall spatial patterns of circuit dynamics parameters we next went on to assess the spatial maps of the *differences* in parameters between CHR-P or FEP groups versus controls and between the drug versus placebo conditions. We computed the difference maps between the average maps for each of the two clinical groups (CHR-P or FEP) versus the HC group. We refer to the parameter difference maps between each of the two clinical groups (CHR-P or FEP) versus the HC group as “psychosis signatures”. Both GABA-A and NMDA receptors (*45*, *60*, *61*) as well as the circuit dysfunctions in schizophrenia (*3*, *29*) are expressed across many cortical areas, but with heterogeneous distribution, possibly over and above the large-scale gradient determined by anatomical hierarchy. We, therefore, hypothesized that the psychosis signatures of the two clinical populations might exhibit characteristic large-scale spatial patterns.

As expected, the psychosis signatures were widespread across cortex (Figure 2). We show the parameter-wise signatures for the aperiodic exponent in Figure 2A and for all other parameters in Figure S2 A-E (same organization in Figure 3). For the aperiodic exponent, the psychosis signature exhibited considerable heterogeneity, in that some areas showed smaller exponents in the clinical group (blue in Figure 2A) while other areas showed larger exponents (red in Figure 2A; similar for other parameters: Figure S2 A-E). Importantly, this heterogeneity in the psychosis signature, while unrelated to anatomical hierarchy (Figure 2B), was highly similar between CHR-P and FEP groups (r(180)=0.85, p=0.0005; Figure 2C). A similar consistency between the two clinical groups was observed for the maps of the disease-related changes in the other parameters (Figure S2 F). This remarkable consistency of the spatial patterns of parameter differences between two independent datasets and groups with different clinical characteristics demonstrates that the psychosis signatures identified by our approach are early and stable markers of the pathophysiology of psychosis that are already detectable in the pre-diagnostic (CHR-P) stage.

**Figure 2:**
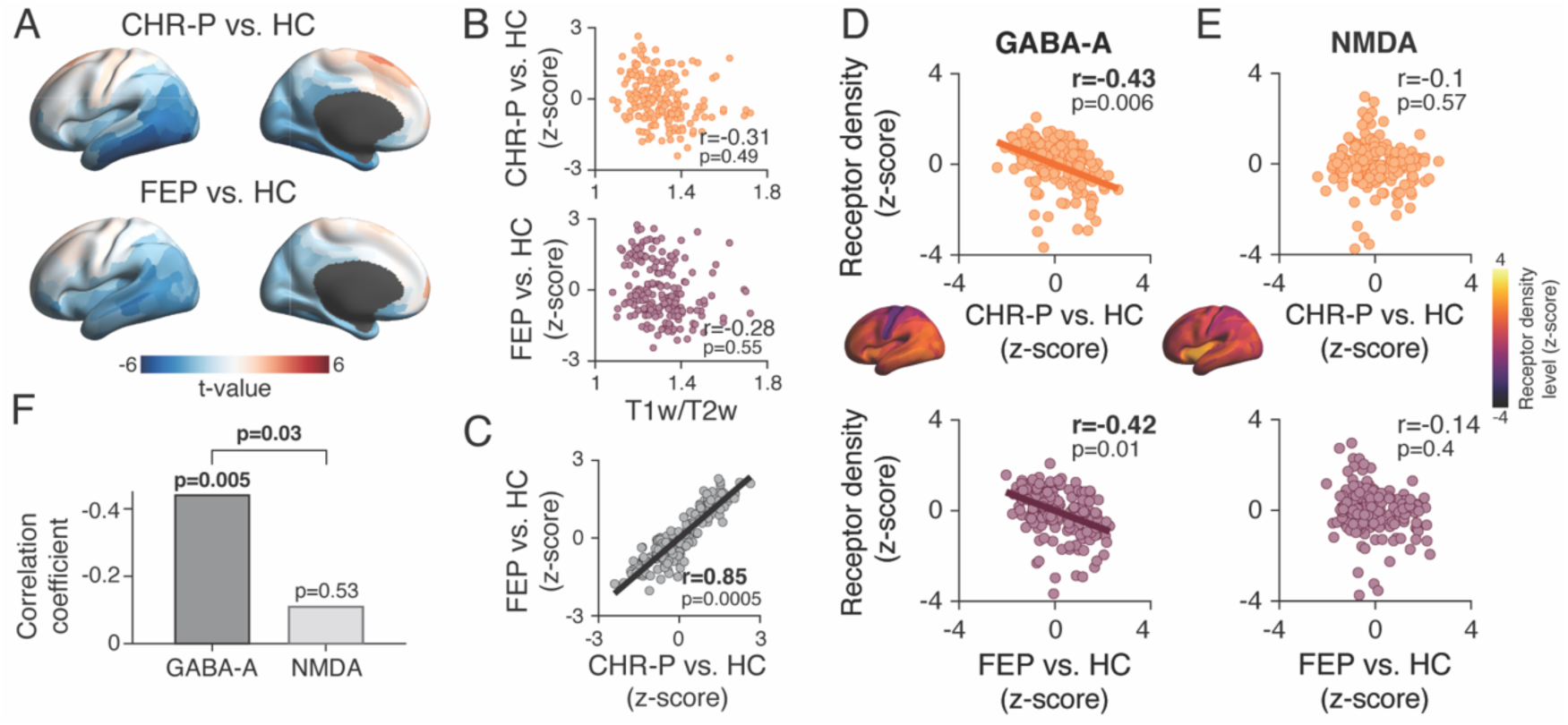
Reproducible large-scale pattern of psychosis signature on exponent of aperiodic activity. **(A)** Cortical distributions of the psychosis signatures (difference map between each clinical group versus HC group) on the aperiodic exponent as group level t-maps. CHR-P: clinical high risk; FEP: first episode psychosis; HC: healthy control. **(B)** Spatial similarity between the maps of psychosis signatures on aperiodic exponent and of T1w/T2w ratios. Each dot represents a single cortical area. **(C)** Spatial similarities of psychosis signatures on aperiodic exponent in the two clinical groups. Black line: linear fit. **(D, E)** Spatial similarity of maps of psychosis signature on aperiodic exponent and maps of GABA-A (D) or NMDA (E) receptor densities (insets), separately for both clinical groups (top/orange: CHR-P; bottom/purple: FEP). Lines: linear fit. **(F)** Differences in similarity between map of psychosis signature on aperiodic exponent (pooled across CHR-P and FEP groups) and GABA-A versus NMDA receptor densities; significance of difference in correlation tested by means of two-sided permutation test. Throughout, r-values are Spearman’s correlation coefficients, p-values are corrected for spatial autocorrelation. In all scatter plots, data points are cortical areas and lines are linear fits for significant correlations.

**Figure 3:**
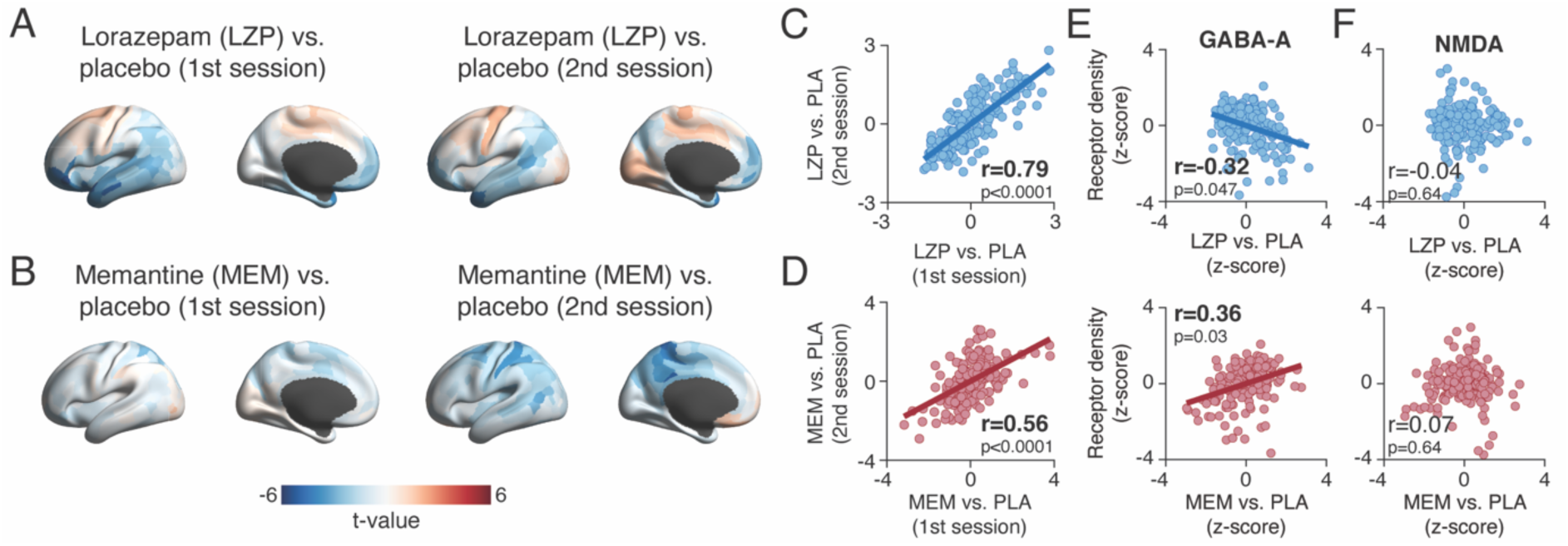
Reproducible large-scale patterns of drug effects on exponent of aperiodic activity. (A,B) Maps of drug effects on the aperiodic exponent (group-level t-maps) for the first (left) or second (right) repetition of each drug condition across different sessions. (A), lorazepam (LZP); (B), memantine (MEM). **(C,D)** Spatial similarity of drug effects in the first versus second administration of each drug condition, a measure of test-retest reliability. (C), LZP; (D), MEM. **(E,F)** Spatial similarity of maps of drug effects on aperiodic exponent and maps of GABA-A (E) or NMDA (F) receptor densities (top/blue: LZP; bottom/red: MEM). Throughout, r-values are Spearman’s correlation coefficients, p-values are corrected for spatial autocorrelation, and lines are linear fits for significant correlations.

The psychosis signatures for the FEP and CHR-P groups also resembled the cortical distributions of GABA-A (not NMDA) receptor densities (Figure 2D-F). We again used spatial correlations to quantify the similarity between the respective psychosis signatures and maps of GABA-A or NMDA receptor densities from a PET study of healthy participants (*45*). The GABA-A receptor density map correlated with the psychosis signature maps for aperiodic exponent, for both clinical groups (Figure 2D). No such relationship was found for the map of NMDA receptor densities (Figure 2E). These pattern correlations differed significantly between both receptor types (Figure 2F).

One possible explanation of the absence of correlation with NMDA receptor density maps may be a smaller across-cortex variation in the latter, compared to the GABA-A receptor density maps. This possibility is not supported by the data, because both receptor maps (insets in Figure 2D, E) had similar statistical properties (spatial variance: GABA-A:0.95, NMDA: 0.93; spatial autocorrelation Moran’s I: GABA-A:0.074, NMDA: 0.06; see Methods). Except for alpha power (GABA-A receptor densities vs. FEP signatures), and alpha peak frequency (NMDA receptor densities vs. CHR-P and FEP signatures), such spatial relation to GABA-A receptor density maps were not evident for the other parameters of circuit dynamics (Figure S3).

In sum, the spatial pattern of the change in the aperiodic exponent in CHR-P and FEP-groups reflects the spatial distribution of GABA-A (but not NMDA) receptors in the healthy brain, pointing to a role of the cortical distribution of GABAergic neuronal inhibition, and the resulting effect on local cortical excitation-inhibition ratios, in early-stage psychosis.

### Relating patterns of changes of circuit dynamics due to psychosis and pharmacological manipulations in healthy participants

We next assessed whether and how the large-scale neurophysiological psychosis signatures identified above related to the changes in cortex-wide dynamics induced by pharmacological manipulations of GABA-A and NMDA receptors in healthy participants. Administration of lorazepam (GABA-A receptor stimulation) and memantine (NMDA receptor blockade) produced widespread changes in the aperiodic exponent compared to placebo. In the following, we refer to the corresponding difference maps as “drug effect maps”. Like the psychosis signatures, the drug effects differed across cortical areas, with smaller exponents (blue in Figure 3A,B) in some areas and larger exponents in other areas (red in Figure 3A,B). Similar heterogeneity was observed for the drug effects on other dynamics parameters (Figure S2 A-E).

These drug effect maps were stable and reproducible features, which reflected the distribution of synaptic properties (Figure 3C-F). The pharmacological study entailed two MEG sessions for each pharmacological condition (separated by 1 to 5 weeks). The spatial patterns of both the effects of both drugs were highly consistent across these two condition repeats for the aperiodic exponent (Figure 3C,D) as well as on most of the other parameters (Figure S2 H). Likewise, the maps of drug effects on exponent (but not the other parameters) exhibited significant spatial similarity to the map of GABA-A receptor densities (Figure 3E), but not the map of NMDA receptor densities (Figure 3F), with opposite signs for lorazepam and memantine effects (Figure 3E). These results establish the reproducibility and significance of the large-scale spatial patterns of changes in cortical dynamics induced by the two drugs.

We used two approaches to quantify the spatial similarity of changes in cortical dynamics associated with psychosis (psychosis signature maps) to those induced by the pharmacological interventions in healthy participants (drug effect maps). The first approach used canonical correlation analysis (CCA), a multivariate statistical technique that determines the linear combinations of two sets of random variables (i.e., matrices) that maximizes the correlation between the two. Here, the random variables were the spatial maps of psychosis signatures or drug effects in the six cortical dynamics parameters defined in Figure 1. We horizontally concatenated these spatial maps, yielding two matrices with 180 rows (areas) and six columns (parameters; Figure 4A). Separate CCAs were performed for the LZP and the MEM effects, respectively. The variables resulting from the linear transformations are called “canonical variables” (Figure 4, B,C). Here, these were again sets of spatial maps (for psychosis signatures and drug effects; Figure 4C), which were ordered by the fraction of co-variance explained by each pair of variables (Figure 4 D,E).

**Figure 4:**
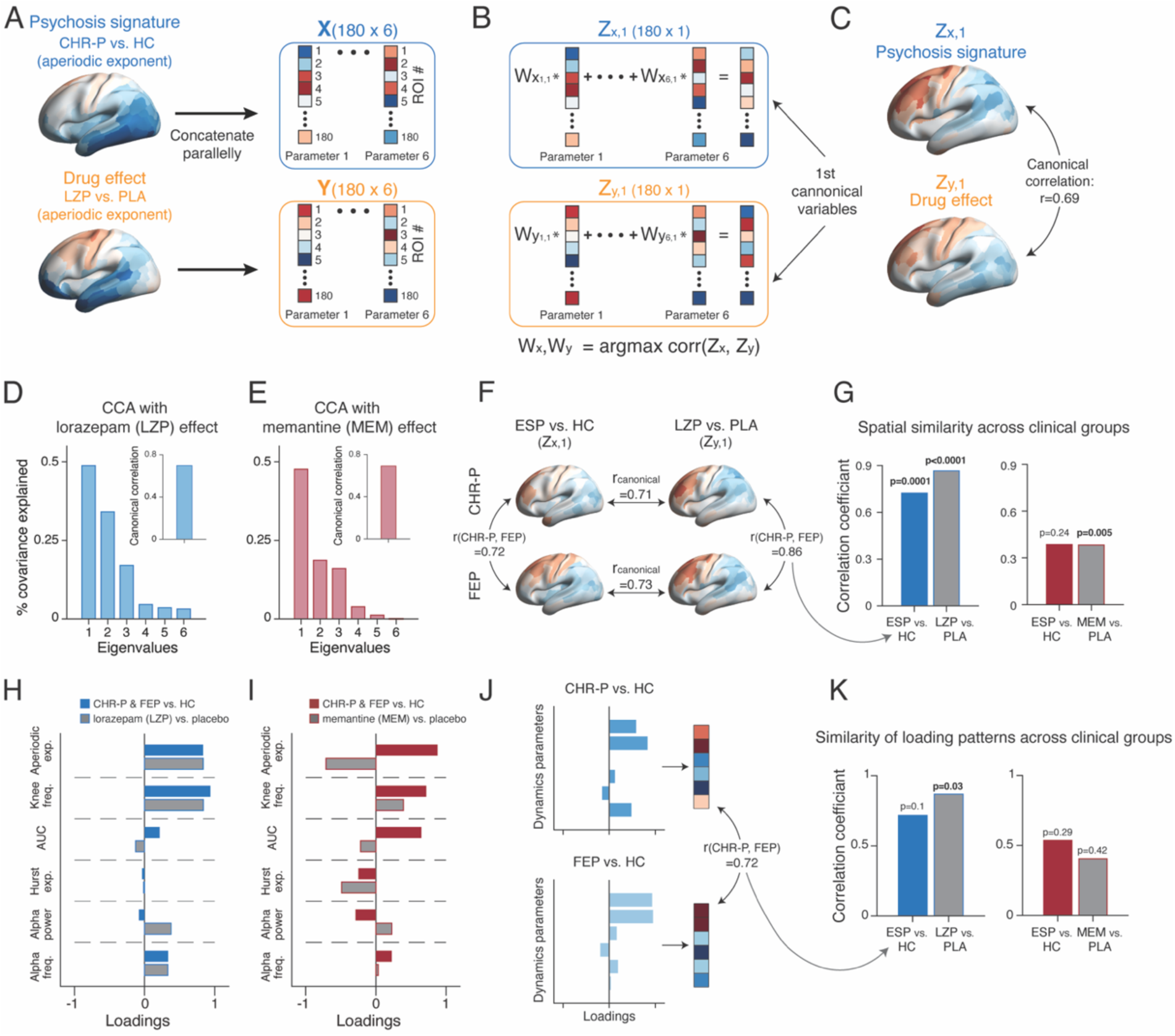
Exponent and knee frequency dominate similarity between psychosis signatures and drug effects. (A-C) Illustration of canonical correlation analysis (CCA) approach. **(A)** Maps of psychosis signatures (e.g., CHR-P vs. HC) or drug effects (e.g., lorazepam vs. placebo) on the six parameters (left) were concatenated parallelly to form two matrices (*X*, *Y*; respectively), each with 180 rows (areas) and 6 columns (parameters; right). **(B)** CC determined matrices (dimensionality: 6x6) of linear weights (*W*_*x*_, *W*_*y*_). Multiplication of these weight matrices with *X* and *Y*, respectively, (*Z*_*x*_ = *XW*_*x*_ *and Z*_*y*_ = *YW*_*y*_) yielded canonical variables (six column vectors of *Z*_*x*_ and *Z*_*y*_) that maximized the spatial correlation. **(C)** Maps of the first canonical variables *Z*_*x*,1_ and *Z*_*y*,1_, respectively. **(D, E)** Spectra of eigenvalues quantifying the fraction of co-variance between *X* and *Y* explained by each pair of canonical variables. Insets: canonical correlation coefficients of the first canonical variables. **(H, I)** CCA loadings reflecting the relative contribution of each parameter to the canonical correlation. **(F)** Maps of the first canonical variables of the CCA between the pooled psychosis signature (Early-Stage Psychosis; ESP vs. HC) and lorazepam effect (LZP vs. PLA). The maps were computed separately for the CHR-P and FEP groups. Horizontal arrows: canonical correlation. Vertical arrow: spatial similarities of corresponding canonical variables across groups. **(G)** Spatial similarity of first canonical variables across the two clinical groups. **(J)** CCA loadings of the first canonical variable between the psychosis signature and lorazepam effect, computed separately for the CHR-P and FEP groups. Horizontal arrow: correlation between loadings of the two clinical groups. **(K)** Similarities between loadings across the two clinical groups. Blue: LZP; red: MEM. P-values were obtained using permutation tests (corrected for spatial autocorrelation in (G)). ESP, early-stage psychosis (pooled psychosis signatures for CHR-P FEP).

The first set of canonical variables accounted for a substantial fraction of the co-variance shared between the psychosis signature and drug effect: 48% for LZP (Figure 4D) and 47% for MEM (Figure 4E), each more than 10% larger than the co-variance explained by the next set of canonical variables. We, therefore, focused on the first canonical variables. These yielded a strong canonical correlation of r(180)=0.69 for both drug effects (Figure 4D,E, inset). Note that only the magnitude, not the sign, of the canonical correlations were meaningful because canonical correlates are positive by construction.

Akin to the psychosis signature (Figure 2) and drug effect (Figure 3) maps, the maps of canonical variables were highly reproducible across independent data sets (Figure 4 F,G; Figure S4 I-L). Consider the similarity of the two canonical variables between repeated sessions for a version of the LZP-CCA (Figure 4F). Correlations between the maps within each row are the corresponding canonical correlations, while the correlations between the canonical variable maps within each column quantify their consistency across the two clinical samples (drug effects pooled across repeats of each drug condition). We ran each CCA separately for the psychosis signature patterns in the CHR-P and FEP groups and separately for the drug effect maps in the two sessions of each drug condition. For each of the LZP -CCAs and MEM-CCAs, this yielded a total of four pairs of first canonical variables (Methods): two pairs of maps for the clinical groups (LZP: Figure S4 A/C and B/D) times two pairs of maps for the session repeats (LZP: Figure S4 A/B and C/D; see Figure S4, E-H for the corresponding results of the MEM-CCA). These maps exhibited substantial spatial similarities (r(180)≥0.28), most (21/24) of which were statistically significant after spatial autocorrelation-preserving permutation tests (Figure S4, I-L).

In the same way, we assessed the consistency of the loading patterns between repeats of CCAs across the two clinical groups and drug sessions (Figure 4 H-K; Figure S4 M-P). The loadings of the canonical variables quantified the relative contribution of the six parameters of cortical dynamics to the canonical correlation. Under lorazepam, the loadings were the largest for the aperiodic exponent and knee frequency, each with equal sign for psychosis signature and LZP effect, (Figure 4H), indicating that the spatial patterns of the changes of both parameters due to lorazepam and psychosis were positively correlated. This supports the visual impression that the maps of changes in exponent in both clinical groups (Figure 2A) and under lorazepam (Figure 3A) resembled one another. By contrast, under memantine, the aperiodic exponent had large loadings of opposite sign between drug effect and psychosis signature (Figure 4I), indicating an anti-correlation between their respective spatial patterns. The loading patterns, too, were fairly consistent across independent data sets. Figure 4K shows their similarity across the two clinical samples (drug effects pooled across repeats of each drug condition), with high and significant correlations for the LZP-CCA (not the MEM-CCA). Many correlations of the loading patterns across the different repeats of CCAs were statistically significant (13/24; 2 further marginally significant at p=0.06), but more consistently for LZP than for MEM (Figure S4 M-P).

Taken together, the CCA approach established substantial large-scale pattern similarity between the psychosis signatures of the CHR-P and FEP-groups on the one hand and drug effects on the other hand. These similarities were reproducible and consistent across two independent clinical groups and experimental sessions separated by several weeks. The approach also uncovered the relative contribution of the six dynamics parameters to this pattern similarity, highlighting the contributions of the exponent and knee frequencies of the aperiodic activity components.

### Individual differences in similarity levels account for clinical symptoms

CCA is a principled approach to relating multivariate sets of spatial patterns, but it also has limitations: it required (i) a focus on the relationships captured by a subset (in our case the first) of the canonical variables, and (ii) averaging the psychosis signatures across the groups of participants. We reasoned that the latter limitation may be important with respect to psychosis, a condition known to exhibit considerable inter-individual variability, at the genetic and molecular levels (*55*) as well as at the level of circuit function (*3*). In other words, different patients diagnosed with, or at risk of developing psychosis might exhibit distinct spatial “fingerprints” of neurotransmitter alterations.

We, therefore, devised a second approach that enabled us to also assess the inter-patient variability in the pattern similarities between their individual effects (compared to the group average HC maps) and the effects of the pharmacological interventions in healthy participants (Figure 5). Here, we concatenated all parameters vertically, yielding a single column vector (*u_scz_*), and for each drug effect (*v_LZP_* and *v_MEM_*). Critically, we computed *u_individual_* as the difference between each individual parameter map from the CHR-P and FEP groups and the corresponding group average parameter maps for the HC group, yielding individual psychosis signatures. Likewise, *v_LZP_* and *v_MEM_* were computed by pooling the data of all healthy participants from the pharmacological study to provide a group-average drug effect as a common reference. We then computed the similarity between psychosis signature and drug effect as the correlation between each individual patient effect and these pharmacological reference patterns (i.e., group average LZP or MEM effect; Figure 5A).

**Figure 5:**
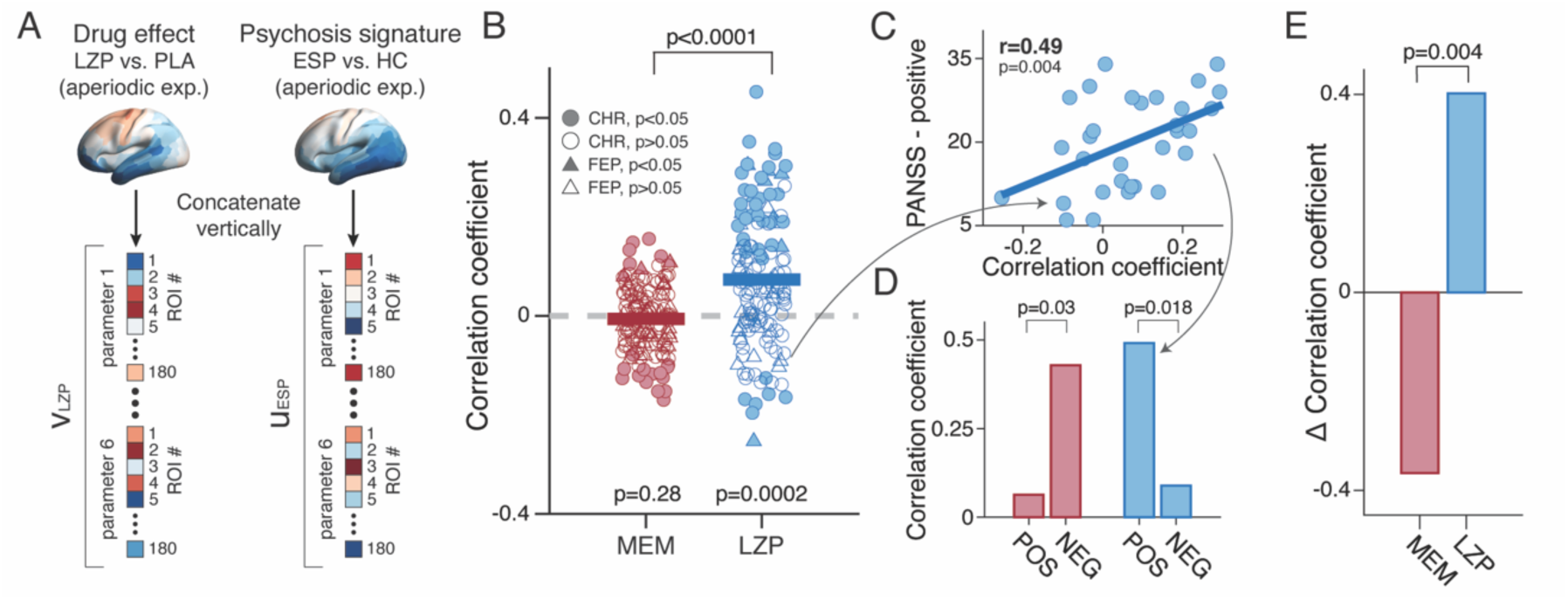
Across-patient variations of pattern similarity to drug effects relates to individual clinical symptoms. **(A)** Illustration of approach for individual difference analysis. Maps of the drug effect or psychosis signature (each of which were column vectors with 180 elements) on the complete set of six parameters were concatenated vertically to form “hyper-vectors” with 1080 elements (6 parameters x 180 areas). **(B)** Spatial similarity between group-average drug effects and individual psychosis signatures (individual patient vs. HC average). Each dot represents the level of pattern similarity for a single patient and a given drug effect (LZP or MEM). Filled symbols, individual patients with statistically significant pattern similarity with a given drug effect. **(C)** Correlation between individual pattern similarity levels from panel (B) and a clinical summary score (so-called PANSS positive scores). The PANSS score was obtained only from FEP patients. **(D)** Correlation between the individual spatial similarities from B and PANSS negative/positive scores, for memantine (red) and lorazepam (blue). **(E)** Comparison of the differences between correlations for positive and negative symptoms between drug conditions. The corresponding p-value tests for the significance of this comparison quantifies a double dissociation (factors: drug and PANNS dimension). All p-values were assessed with two-sided permutation tests (corrected for spatial autocorrelation in (B)).

This approach revealed substantial variability across CHR-P and FEP individuals in the similarity levels between their psychosis signatures and the group-average drug effect (Figure 5B; Figure S5). For both LZP and MEM reference patterns, we found statistically significant similarities (correlations) for individuals from both clinical groups of opposite signs (Figure 5B; filled symbols above and below 0). For the MEM reference pattern, a comparable number of patients exhibited positive or negative correlations. When pooling the individual correlations across the CHR-P and FEP groups (based on their high similarity, Figure 2C), the group average correlation coefficient that was indistinguishable from zero (r(1080)=-0.006; p=0.28; Fig. 5B; see Figure S5 for separate tests for each clinical group). The individual LZP similarities were mostly positive and yielding a significantly positive group average correlation (pooled across CHR-P and FEP: r(1080)=0.07; p=0.002, Fig. 5B; Figure S5).

This large variability in similarity levels was related to patients’ individual clinical symptoms (Figure 5C-E). In the FEP group, clinical symptoms were assessed in terms of the Positive and Negative Symptoms Scale (PANSS; (*62*)). We correlated the individual similarities for each drug pattern from Fig. 5B with either the positive or negative aggregate PANSS scores in the FEP-group. This revealed distinct symptom-correlates of the spatial similarities to the LZP and MEM effects: MEM similarities correlated with negative symptoms (r(32)=0.45, p=0.007) while LZP similarities correlated with positive symptoms (r(32)=0.5, p=0.004). The differences in correlation with positive and negative scores for MEM and LZP, respectively, was highly significant effect (p =0.004; Figure 5E), demonstrating the specificity of these effects.

In other words, FEP patients whose spatial patterns of alteration in cortical dynamics exhibited larger similarity to the changes by a boost of GABA-A transmission exhibited more severe positive symptoms, and patients with stronger similarity to changes induced by NMDA-R blockade exhibited more severe negative symptoms.

### No robust local expression of psychosis signatures or drug effects in cortical dynamics

Our approach focused on the similarity of the cortex-wide spatial patterns of the changes in cortical dynamics induced by early-stage psychosis and the two pharmacological manipulations. This focus was motivated from the emerging understanding of the principles of receptor expression in the healthy brain (*39*) as well as of circuit aberrations (*3*) and was corroborated by the observations from Figures 2 and 3, that the psychosis signatures and drug effects in cortical dynamics were widespread, with distinctive spatial patterns across cortex.

Control analyses showed that the local changes in the parameters, irrespective of their large-scale spatial patterns, were not sufficiently strong to be reliably detectable, neither of the clinical disorder nor the pharmacological interventions (Figures 6 and S6). We reduced the number of multiple comparisons (i.e., boost statistical sensitivity) by pooling each local dynamics parameter across areas, and then comparing the resulting aggregate parameters between clinical groups or pharmacological conditions. We pooled at two levels of granularity: (i) across all parcels belonging to each of 22 groups of areas (*44*) (psychosis signatures on aperiodic exponent: Figure 6A; drug effects on aperiodic exponent: Figure 6H; see Figure S5 for all other parameters); and (ii) across the complete set of 180 areas, yielding a single number per parameter (psychosis signatures: Figure 6 B-G; drug effects: Figure 6 I-N). Despite using a lenient threshold (p<0.05), we found few significant effects (Figures 6 and S6). Especially the clinical datasets which yielded only one single significant effect (in the paracentral lobular and mid cingulate cortex (*44*) for the area under the aperiodic curve). Bayes factors provided support for the null hypothesis (i.e., *BF*_12_<1/3 (*63*)) for most comparisons in the clinical dataset (HC vs. CHR-P or FEP; AUC: *BF*_12_<0.27; Hurst exponent: *BF*_12_= 0.19 (CHR-P only); Alpha power: *BF*_12_<0.25; Alpha frequency: *BF*_12_<0.24). These control analyses establish that our large-scale pattern analyses were in fact essential to identify robust psychosis signatures and identify their relationship to drug effects.

**Figure 6:**
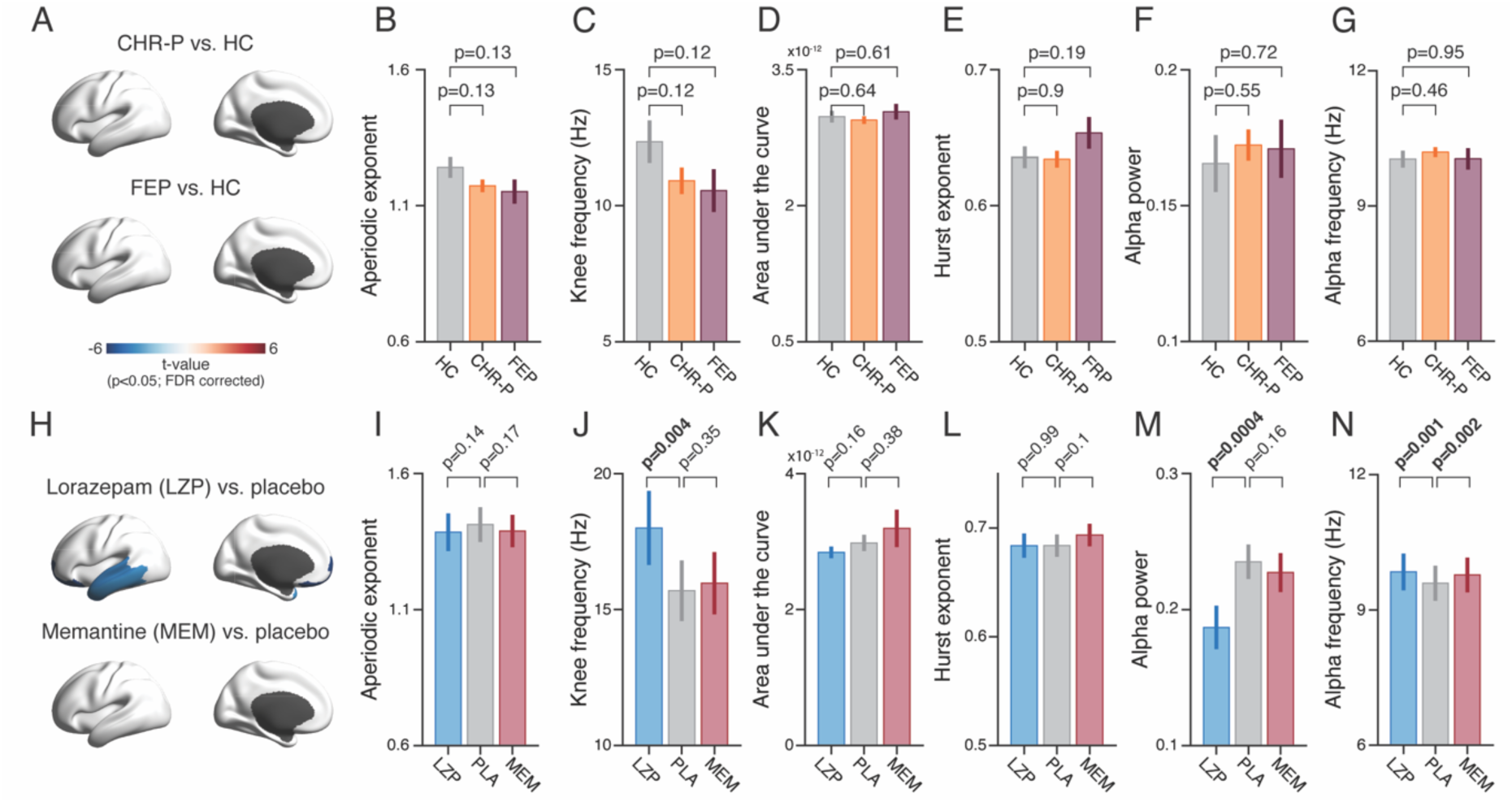
Weak psychosis signatures or drug effects in local cortical dynamics. **(A)** Spatial patterns of the psychosis signatures (CHR-P vs. HC or FEP vs. HC) on the aperiodic exponent, presented as group level t-value of significant parcels only (p<0.05, FDR correction). **(B-G)** Means and standard errors across all 180 parcels for healthy control (HC; grey), clinical high risk (CHR-P; purple) and first episode psychosis (FEP; orange) groups of the aperiodic exponent (B), knee frequency (C), area-under-the-curve (D), Hurst exponent (E), alpha power (F), and alpha peak frequency (G). **(H-N)** same as (A-G) for the pharmacological conditions; LZP: lorazepam (blue); PLA: placebo (grey); MEM: memantine (red). All p-values were assessed with two-sided permutation tests.

## Discussion

The cerebral cortex constantly generates a rich repertoire of dynamical activity patterns, such as band-limited oscillations and aperiodic activity, which have been suggested as potential biomarkers for neuropsychiatric disorders (*9*, *28*, *29*, *33*, *35*, *64*). Recent work from computational neuroanatomy into the large-scale distribution of cortical microcircuit properties, such as cytoarchitectural type or receptor gene expression, identified substantial spatial heterogeneity in these parameters across cortical areas and cortical hierarchy as one overarching organizing principle (*22*, *39*). Here, we present an approach to investigate the pathophysiology of psychosis that integrates these different bodies of work: we assessed large-scale cortical patterns of early-stage psychosis-induced changes in several parameters of cortical dynamics and related these patterns to the patterns of changes in the same parameters induced by pharmacological manipulations of GABA-A or NMDA receptors in the healthy brain. In doing so, we leveraged the high spatio-temporal resolution and cortex-wide spatial coverage afforded by source-level analyses of MEG data (*48*), combined with state-of-the-art anatomical atlases (*44*).

Our analyses revealed that the large-scale patterns of alterations in cortical circuit dynamics in psychosis (i) resemble the large-scale patterns of changes in the same dynamics parameters due to pharmacological manipulation of GABA-A receptors, in a way that (ii) reflects the densities of GABA-A receptors across cortical areas and (iii) predicts the individual expression of (positive, but not negative) symptoms. Weaker similarities across both clinical groups were identified for changes in the parameters of cortical dynamics due to manipulation of NMDA receptors. But again, the individual similarity levels were related to the individual expression of symptoms, only in this case the negative symptoms, but not the positive symptoms. It is noteworthy that we found weaker evidence for group-average similarity of psychosis signatures to the pattern induced by NMDA-blockade, in the two analyses approaches from Figures 4 and 5, but substantial inter-patient variability in these similarity levels, which was partly related to negative symptomatology. Together, our results indicate that both neurotransmitter systems are involved in the pathophysiology of psychotic disorders (*65*), but contribute in distinct ways to positive and negative symptoms.

Our approach is inspired by insights from multivariate pattern analysis approaches in neuroimaging (*66*, *67*). Conventional univariate (“activation-based”) approaches were designed to detect differences in the activity of individual sources (e.g., fMRI voxels) between conditions. Such local differences, while robust for coarsely defined conditions (e.g., task versus baseline, or high versus low cognitive load), are often too subtle for reliably discriminating between conditions differing only in specific stimulus or cognitive variables (*66*, *67*). These small differences (“biases”) that may exist at the level of single voxels are aggravated by the severe multiple-comparisons problem associated with mass-univariate statistical tests (*67*). By combining the information contained in many such small (but systematic) single-voxel biases, multivariate pattern analyses boost sensitivity and reduce the multiple-comparisons problem, thereby successfully decoding specific mental content (*68*, *69*). Simpler spatial pattern correlations to ours (just at the level of within-area voxel patterns as opposed to large-scale, across-area patterns) were used from the conception of such multivariate fMRI approaches (*70*) and lie at the heart of state-of-the-art representational similarity analyses (*71*). While guided by the same methodological insights and principles, our current approach is designed for a different purpose: identifying reliable large-scale patterns of changes in cortical dynamics induced by pharmacological interventions or brain disorders, and explicitly relating these patterns to one another.

Indeed, we found only weak local differences in cortical dynamics parameters under either pharmacological manipulation of GABA-A and NMDA receptors or in clinical groups compared to healthy controls. The strengths of pharmacological effects will scale with the drug doses, which were low in our study so as to minimize secondary and side effects and maximize specificity of effects for the receptors with highest affinity. However, the psychosis signatures are unlikely to change substantially between different patient samples. This highlights the importance of our approach focusing on spatial pattern similarity for picking up locally subtle, but spatially highly consistent, reproducible, and clinically relevant features of large-scale cortical dynamics. The observations that the large-scale patterns of these psychosis signatures and drug effects were (i) highly reproducible across independent datasets and (ii) reflected important synaptic features of brain organization (distribution of GABA-A receptor density) establish that these patterns are genuine and meaningful.

Several lines of evidence implicate GABAergic cortical inhibition in psychotic disorders (*6*). Patients with schizophrenia show reduced GABA-levels in magnetic resonance spectroscopy (7, 8) and lower concentrations of molecules involved in the transport and metabolism of GABA in postmortem assays (6, 72, 73). Our findings are overall consistent with this, and they may appear less consistent with body of evidence implicating glutamatergic deficits and NMDA receptors in psychotic disorders (9, 10, 28, 38), in particular early-stage psychosis (REF) (74). In our analyses, the *patient-average* pattern of psychosis signature on cortical dynamics was neither related to the distribution of NMDA receptor density levels, nor to the corresponding of memantine effects in healthy participants. One possible explanation, to be tested in future work, is that the alterations in circuit dynamics due to NMDA hypofunction in psychosis are more prominently expressed during effortful cognitive processes that involve synaptic reverberation, such as working memory or evidence accumulation (10, 19, 38).

Ketamine, another NMDA receptor antagonist, has strong effects on the aperiodic exponent at strong dosage (75) (but not at lower dosage (76)) and, at subanesthetic levels, can produce symptoms in healthy participants that resemble symptoms of psychosis (77–80). Memantine effects on cortical dynamics (specifically, aperiodic exponent) also increases with dosage (81). We used memantine instead of ketamine and a low dosage because (i) memantine and ketamine have analogous mechanisms of action on NMDA receptors (82), (ii) the oral administration of memantine matches the one of lorazepam, improving the blinding in our within-participants intervention, and (iii) low dosage help to minimize non-specific secondary effects (82). Despite the small group average effects, our spatial similarity analyses identified large differences in the levels of similarity between memantine effects and the psychosis signatures expressed in individual early-stage psychosis patients, with highly significant but oppositely signed spatial correlations in different patients, which account for the small patient-average effects. Critically, these inter-individual differences in similarity levels explained a substantial fraction of inter-individual differences in negative symptoms, highlighting the potential of our spatial similarity approach for individualized approaches in psychiatry.

The similarity between GABA-A receptor density and both, average psychosis signatures and drug effects were most evident in the aperiodic exponent. This parameter is sensitive to changes the excitation-inhibition ratio in local circuits (33), and increases in the aperiodic exponent (averaged across all electrodes) have been reported in schizophrenia (81). Critically, however, the aperiodic exponent did not suffice to explain the relationship between similarity levels of individual patients with drug effects and positive or negative symptoms: repeating our pattern analyses from Figure 5 using only the aperiodic exponent did not show similarly clear effects. This indicates the importance of pooling the information provided by several distinct features of cortical dynamics for detecting the neurophysiological alterations underlying psychotic disorders.

A large body of work has demonstrated psychosis-related aberrations of neuronal responses in the gamma-band (14, 29), which are, in turn, generated by excitatory-inhibitory synaptic interactions within cortical microcircuits (57–59, 83). We here focused on alpha-rather than gamma-band oscillations for the quantification of periodic components of local cortical dynamics. This choice was pragmatic, because our approach relied on across-area patterns, and alpha peaks were more consistently detected across cortical areas in the resting-state data analyzed here. Under conditions of strong external (stimulus and/or task) drive of local cortical circuits, gamma-band responses will be more prominent (58), and future applications of our approach to such data may also utilize gamma-band oscillations. That said, the alpha-band oscillations used here are likely to provide relevant information complementary to the information caried by parameters describing the aperiodic activity components, for example due to a contribution of thalamocortical interactions to the generation of cortical alpha-band oscillations (84).

We replicated the observation from previous work (34, 42, 85, 86), that the spatial patterns of cortical dynamics parameters followed the anatomical hierarchy (39). Importantly, we here show that this relationship is preserved in early-stage psychosis as well as under pharmacological manipulations of NMDA and GABA-A receptors. Anatomical hierarchy has been identified as an important principle that accounts for heterogeneity in microcircuit architecture and the dominant axis of gene-expression variation across the cortex (22, 39). By contrast, we here show here that the psychosis signatures and drug effects in the spatial patterns of dynamics parameters were only weakly associated with anatomical hierarchy, but strongly associated with GABA-A receptor density levels. This points to the presence of other principles of large-scale brain organization beyond anatomical hierarchy (87, 88), which may be important for understanding the pathomechanisms of psychotic disorders.

We conclude that early-stage psychosis is associated with consistent large-scale patterns of changes in multiple parameters of cortical dynamics, referred to as psychosis signatures, which (in part) result from alterations in GABA-A or NMDA receptor functions. Individual differences in those psychosis signatures relate to individual symptomatology. Our results are consistent with the notion that mechanisms of psychosis are fundamentally distributed across many regions of the cerebral cortex. The approach we developed opens up perspectives for a mechanistically informed stratification of patient cohorts in future psychosis research.

## Methods

We analyzed resting-state MEG data sets collected in two separate studies: (i) a study (50) comparing CHR-P participants, FEPs versus healthy controls; and (ii) a pharmacological study(51) in young and healthy participants with double-blind and placebo-controlled manipulations of GABA-A or NMDA receptors, respectively.

### Participants and informed consent

#### Clinical dataset

Three groups of participants were recruited to the study. One group included participants who met CHR-P criteria (N=117, mean age: 22 y, range: 16-24 y, 83 females), based on the Comprehensive Assessment of At-Risk Mental State interview (CAARMS) (89), and the Cognitive-Perspective Basic Symptoms (COGDIS/COPPER) item of the Schizophrenia Proneness Instrument (SPI-A) (90). Exclusion criteria included diagnosis of Axis I psychotic disorders, including affective psychosis, as evaluated by the Structured Clinical Interview for the Diagnostic and Statistical Manual of Mental Disorders-IV (SCID). In addition, an FEP-group (N=32, mean age: 24 y, range: 18-34 y, 12 females) was recruited. Psychopathological symptoms were evaluated using the Positive and Negative Symptoms Scale (PANSS) (62).

The control group for both above clinical groups was a sample of age- and gender-matched healthy controls (HC; N=45, mean age = 23 y, range: 18-32 y, 31 females), who were screened for psychopathology using the SCID or the MINI-SCIS interview. The HC group was matched by age to the two other group (HC vs. FEP and CHR-P combined: p=0.48, two-sided permutation test), but not by years of education (HC: 16.8 y; CHR-P and FEP combined: 15.1 y; p=0.002, two-sided permutation test). There was a similar percentage of females in the HC and in the combined clinical sample (68.8% and 63%, respectively; p=0.47; Chi-square test). The three groups of participants were recruited as part of the Youth Mental Health Risk and Resilience (YouR) study (see (50) for more information about subjects’ recruitment, exclusion criteria and experimental procedures), which was approved by the NHS Research Ethical Committee Glasgow. All participants provided written informed consent and received remuneration of 36 £. We excluded 3 participants (all from the CHR-P group) due to excessive artifacts in MEG signals.

#### Pharmacological dataset

We recruited 23 healthy participants (mean age: 28 y, range: 21-40 y; 9 females) to the study. Exclusion criteria included: impaired vision or hearing, illegal drug use, regular medication intake, consumption of more than 15 units of alcohol per week, known hypersensitivity to Memantine or Lorazepam, past or current psychiatric or neurological diagnosis, cardiovascular, liver, kidney or metabolic diseases, pregnancy, and non-removable metallic parts (e.g., insulin pump, retainer). We excluded one participant after detecting excessive metal artifacts in the MEG signal, and two participants dropped out before completing all experimental sessions. Participants received remuneration of 15€ for the training session, 100€ for each MEG session, 150€ as a completion bonus, 10€ for MRI session, and a performance-dependent bonus of a maximum of 150€. The study was approved by the ethics committee of the Hamburg Medical Association, and all participants provided written informed consent.

### Experimental design

#### Clinical dataset

The three groups of participants (CHR-P, FEP and matched controls) completed one MEG session (∼ 3.5 h long) followed by one MRS/MRI session (∼2.5 h). Among other tasks in the MEG (50, 91), participants completed a 5 min block of resting state recording with eyes open (fixation of a central cross on an otherwise gray screen). Participants were instructed to keep fixation at a plus sign in the center of a gray screen and keep a blank state of mind as well as possible, (i.e., do not think of anything in particular and if a thought comes up, do not follow up on it).

#### Pharmacological dataset

Participants completed one training session (outside the MEG) followed by six MEG sessions, interleaved by at least one week. Each MEG session started with the intake of a pill (Lorazepam, Memantine or placebo; see ‘Pharmacological manipulation’ subsection), followed by a 150 min waiting period before MEG recording. In the MEG, participants performed several different tasks, lasting approximately 2.5 h. This included a 10 min block of resting state recording with eyes open (fixation of a central cross on an otherwise gray screen). This resting state recoding took place at the end of the session, around 4.5 h after drug intake. During the block, participants seated quietly with their eyes open, and asked to maintain fixation at a centrally presented black cross against a grey background throughout the block.

We orally administered lorazepam or memantine, in a double blind, placebo-controlled and a crossover design. Each drug was given twice, in two different experimental sessions, in addition to two placebo sessions. Lorazepam is a GABA-A receptor agonist that boosts GABAergic neurotransmitters. Memantine is a NMDA receptor antagonist that decreases glutamatergic neurotransmitter. We choose sub-clinically dosage for the drugs: 1 mg for lorazepam (common clinical daily use between 0.5 and 2mg) and 15 mg for memantine (common clinical dose: 20 mg). Peak plasma concentration for lorazepam is 2-3 h, and 3 to 8 h for memantine. Thus, we administered the drugs 3 h before MEG session started (2.5 h waiting period plus 30 min preparation time before MEG recording), to maximize plasma concentration levels. To allow plasma concentration level to return to normal, experimental sessions were interleaved by at least one week (plasma half-life of lorazepam is ∼13 h, and of memantine is 60-70 h).

### Data acquisition

#### Clinical data

*MEG:* MEG data was collected at the Center for Cognitive Neuroscience at University of Glasgow, using 248 channels 4D-BTI system (4D-Neuroimaging, San Diego), at a sampling rate of 1017.25Hz. Online low-pass filter was applied at 400Hz.

*MRI:* T1-Weighted data was obtained from 178 out of 191 participants included in the final analyses reported here, using a 3T Siemens Magnetom Trio MRI scanner at the Center for Cognitive Neuroscience at University of Glasgow with the following parameters: 192 slices, voxel size = 1x1x1 mm^3^, FoV= 256x256x176 mm^3^, TR=2250 ms, TE=2.6 ms, flip angle = 9°.

#### Pharmacological data

*MEG:* MEG data was collected at the Department of Neurophysiology at University Medical Center Hamburg-Eppendorf using a CTF MEG system with 275 gradiometers, and a sampling rate of 1200Hz. Participants were seated in a dark and magnetically shielded room and were asked to minimize head movements. We monitored and recorded head positions continuously, using three fiducial coils, placed on participants’ ear canals and nasal bridge. Additionally, we recorded ECG, and vertical and horizontal EOG using Ag-AgCl bipolar electrodes with a sampling rate of 1200Hz. Eye movements were recorded using EyeLink 1000 system, with a sampling rate of 1000Hz. We calibrated gaze positions at the beginning of the block.

*MRI:* T1-weighted images were obtained from all participants using a 3T Siemens Magnetom Trio MRI scanner (Siemens Medical Systems, Erlangen, Germany) and the following parameters: voxel size = 1x1x1 mm^3^, TR=2300 ms, TE=2.98 ms, FoV= 256 mm, slice thickness = 1mm, TI= 1100 ms, flip angle = 9°.

### MEG data analysis

We analyzed both datasets (pharmacological and clinical) using the same procedures, with several adjustments due to differences in the acquisition protocols. Unless stated otherwise, the following refers to both datasets. We analyzed MEG data using customized code from Fieldtrip toolbox(92) in MATLAB (MathWorks), and MNE(93) and pymeg (https://github.com/DonnerLab/pymeg) toolboxes in Python. We preprocessed MRI scans using FreeSurfer.

*Preprocessing.* Preprocessing of MEG data included three steps. The first step detected MEG artifacts to be rejected from the signal before applying independent component analysis (ICA; second step), to avoid extreme values affecting the ICA. The third step rejected data segments containing previously identified artifacts and removed ICA components from the remaining continuous data. Each step applied different high-pass filters with cutoffs adapted the respective purpose. Hence, each step started from the raw unfiltered data.

The first step identified segments containing head movements, muscle contractions, squid jumps, or metal artifacts. Continuous MEG and external electrodes (ECG) were down-sampled to 400Hz, and band-pass filtered to remove powerline noise at 50 Hz, 100 Hz and 150 Hz (two pass Butterworth filter; band stop frequencies: ±1Hz around each frequency). We detected muscle artifacts by band-pass filtering MEG signals between 110Hz to 140 Hz and z-scoring; values exceeding z=10 were labeled as muscle artifacts. Head movements (only for pharmacological dataset; continuous monitoring of head movements was not done in the clinical dataset) were detected as deviation of any coil more than 6mm from the template head position. Squid jumps were detected by fitting a line to the log power spectrum of individual artificial trials (7 s long) and detected outliers of its intercept. Finally, metal artifacts (e.g., cars passing near the building), were detected as samples exceeding a threshold of ± 5 pT.

In a second step, the raw MEG data was high-pass filtered at 1 Hz, and segments containing squid jumps, head movements and metal artifacts were rejected, to avoid extreme values during ICA. We then applied ICA using the infomax algorithm as implemented in MATLAB) and manually identified components containing eye movements (i.e., blinks and saccades) and heart beats.

In a third step, we resampled the raw data and removed power line noise, using the same parameters described above for the first step, high-pass filtered the data at 0.1 Hz (Butterworth filter) and removed previously identified ICA components or rejected data segments previously identified as containing artifacts (muscle contractions, head movements, metal artifacts and squid jumps), respectively. We did not reject segments containing eye movement-related activity (blinks or saccades), but rather removed ICA components containing these artefacts. This was done to match the pre-processing steps between the two datasets, where the clinical dataset did not include any monitoring of eye-movements (e.g., neither EOG nor eye-tracking data).

In total, our procedure led to the rejection of 13% of data segments in the clinical dataset, and 26% of data segments in the pharmacological dataset, resulting in time series with duration of 262 sec ± 36 sec (mean and s.d across subjects) or 437 sec ± 56 sec (mean and s.d across subjects), respectively.

*Source reconstruction.* We used linearly constrained minimum variance (LCMV) beamforming to project broadband sensor-level signals into the cortical source space. We constructed a three-layer head model (inner skull, outer skull and skin) for each subject based on their individual MRI scan. In cases where an MRI scan was missing, we used the FreeSurfer fsaverage to reconstruct a template head model. We aligned the head model to MEG data by computing a transformation matrix, using the mean coils positions across the resting state block (for pharmacological dataset), or a fixed coils positions measured at the beginning of the block (for clinical dataset) and the corresponding locations of fiducial coils on the individual head model. We then used the individual MRI scan and FreeSurfer to reconstruct cortical surfaces and parcellate the cortex into 360 parcels, using the anatomical atlas from Glasser and colleagues (44). Based on the head model, we generated a forward model (“leadfields”) for the computation of LCMV filters, that was confined to the cortical sheet (4098 vertices per hemisphere, recursively subdivided octahedron). For each vertex, a set of LCMV filters (one per cardinal orientation) were computed by combining the leadfield matrices of that vertex with a covariance matrix of the sensor-level signal. For each vertex, we chose the source orientation with the maximum source power using LCMV filters. We projected the time series through the LCMV filters into source space and collapsed the data across all vertices within each parcel. This resulted in 360 time series, one for each parcel, of source-level broadband data.

*Quantification of local cortical circuit dynamics.* We estimated power spectral density (PSD) by dividing the time series of each parcel into 10sec epochs, with 50% overlap, transforming each epoch into the frequency domain using the Fourier transform, and averaging the absolute values of Fourier coefficients across all epochs. This resulted in PSDs with a frequency resolution of 0.1 Hz. We smoothed the PSD at 50Hz, 100Hz and 150Hz using linear interpolation (±2 Hz around each frequency) since activity at those frequencies was filtered during preprocessing. We fitted the FOOOF model (52) to each PSD in a range between 1-65 Hz, to avoid the noise floor at higher frequencies (94), and included a knee parameter, based on a visual inspection of the PSDs. If the algorithm failed to fit the model or returned a negative knee frequency, we re-fitted the model without the knee parameter. Additional fitting parameters were as follows: peak threshold: 2 standard deviations, peak limit: [2 16] Hz. We extracted from the model the aperiodic exponent, knee frequency (if fitted, otherwise a NaN was assigned), alpha power and its peak frequency. We noticed that in some cases multiple peaks within the frequency range from 7 to 13 Hz were detected by the algorithm. To identify alpha peak more accurately, we constrained the range of searching by first fitting the model to the mean PSD across V1 and early visual cortex (V2-V4), where alpha activity is the strongest and extracted a single alpha peak frequency and its bandwidth, after visually verifying it is indeed in the alpha range. We used this frequency and its bandwidth (± 2 Hz) to constrain the range of searching for alpha peaks in all other parcels. If still more than one peak was identified within this range, we chose the one with the highest power. In rare cases, no alpha peak could be detected from the visual cortex. In such events, we selected the peak with the highest power within the traditional alpha range of 7-13 Hz. If no peaks were detected whatsoever, we treated those as missing values (i.e., NaNs) for all further analysis. We computed the area under the curve from the aperiodic fit of the PSD, by summing the power between 1-65 Hz. The Hurst exponent of spontaneous fluctuations of amplitude envelopes in the alpha-band oscillations was computed as in (35). In brief, we used Detrended Fluctuation Analysis (DFA), as implemented in NBT toolbox (Neurophysiological Biomarker Toolbox: https://github.com/NBT-Analytics/NBTpublic), with the following parameters: filter range: 8-13Hz (order 100 FIR filter), calculating range: 1-120 s (50% overlap) and fitting range: 3-50 s. Importantly, the six dynamics parameters extracted here were not correlated by construction. Such trivial correlations exist, for example, for the exponent and intercept of the aperiodic component. Our parameter selection ensured that any (across-area) correlations between parameters identified in the data were meaningful and interpretable in terms of underlying mechanism or shared organizational principles (such as anatomical hierarchy).

### Receptor density maps

We used the publicly available volumetric PET images from (45). We used the group averaged PET images of two neurotransmitter receptors: GABA-A and NMDA. PET images were registered into the MNI template and then parceled into 360 parcels according to the above-mentioned parcellation (44), and then z-scored across parcels.

### Canonical correlation analysis (CCA)

We used CCA to relate the psychosis signature on the cortical patterns of the six dynamics parameters to the matching drug effect on them. In the clinical dataset, we first z-scored the cortical spatial pattern of each of the six parameters, computed the mean across both clinical groups (i.e., pooled across CHR-P & FEP), and finally calculated the psychosis signature by subtracting from the mean clinical subjects the mean across healthy controls. Similarly, in the pharmacological dataset, we z-scored each parameter for each subject and drug condition, and then calculated the drug effect as the mean across subjects of the difference between each drug condition and the placebo condition. We then horizontally concatenated the group-average spatial patterns of all six parameters into two 180x6 matrices, which we denoted X for the psychosis signature and Y for the drug effect. We performed two separate CCA’s for the lorazepam and memantine effects, respectively, so below Y refers indiscriminately to lorazepam or memantine effects, since the computation was identical for both effects.

CCA decomposed the relationship between the two multivariate datasets (i.e., the pair of matrices X and Y) into orthogonal sets of latent variables according to:

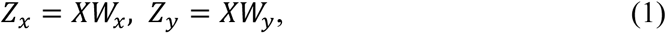

where *W*_*x*_ and *W*_*y*_ were 6x6 matrices of linear weights, which *X* and *Y* were projected onto, resulting in 180x6 matrices *Z*_*x*_ and *Z*_*y*_, the columns of which corresponded to the six latent (so-called “canonical”) variables. The weights were chosen so as to maximize the correlations between each pair of canonical variables (i.e., corresponding columns of *Z*_*x*,_, *Z*_*y*_). These pairs of columns of *Z*_*x*_and *Z_Y_* were ordered by the magnitude of their respective correlations, such that the first pair had the largest correlation, and each subsequent pair had progressively smaller correlations, while being orthogonal to all other pairs. The weight matrices (*W*_*x*_, *W*_*y*_) satisfying these constraints were found by first applying a QR decomposition to X and Y:

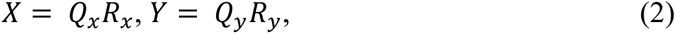

where Q were 180x6 orthogonal matrices (columns were orthogonal unit vectors) of dimensionality, and R were 6x6 upper triangular matrices. The term *Q*_*x*_^*T*^*Q_y_* was subjected to singular value decomposition (SVD):

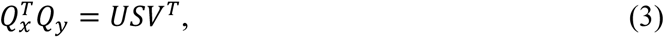

where U and V were 6x6 orthogonal matrices, and S was a 6x6 diagonal matrix containing the canonical correlation coefficients in ascending order, such that *diag*(*S*) = *R*, where R was a vector of canonical correlation coefficients. The weight matrices were computed as:

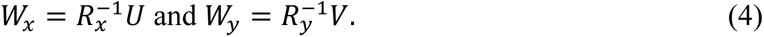

The amount of co-variance explained by each set of canonical variables equalled R^2^.

To quantify the relative contribution of each dynamics parameter to the canonical correlation for the first canonical variables (first columns of *Z*_*x*_ and *Z*_*y*_), we computed the so-called “loadings” as the Pearson’s correlation coefficients between these canonical variables and the original data (psychosis signature or drug effect) matrices. For each CCA (i.e., the LZP and the MEM analysis), this yielded a pair of 1x6 vectors, with one entry per dynamics parameter, for the contributions of parameters in drug effect and in psychosis signature, which were tested further for their reliability as described below.

To demonstrate the reproducibility of the CCA results, we repeated the analysis for each clinical group (i.e., CHR-P and FEP) and for each pair of the repeated sessions (S) of a pharmacological condition (e.g.., LZP S1 and LZP S2). This resulted in a total of 4 pairs for each drug condition: session S1-CHR-P, S2-CHR-P, S1-FEP and S2-FEP. For each combination, we computed the canonical variables matrices (e.g., *Z*_*x*(*CHR*,*S*1)_, *Z*_*y*(*CHR*,*S*1)_) and corresponding loading vectors, and quantified the similarity in the resulting (spatial or parameter) patterns across sessions and clinical groups by calculating the correlation between pairs of canonical variables (e.g., *Z*_*x*(*CHR*,*S*1)_, *Z*_*x*(*CHR*,*S*2)_) or between pairs of corresponding loading vectors, across all combinations. This was done separately for variables or loadings associated with psychosis signatures and drug effects.

### Individual correlations of spatial patterns of effects

We tested the similarity between the effect of memantine or lorazepam on the spatial patterns of cortical dynamics parameters to the individual effect of patients. Parameters were z-scored as described in the section above, and the drug effect was computed in the same way. The individual psychosis signature, however, was computed as the difference between each patient and the mean across all subjects in the healthy control group. We then concatenated all parameters vertically into a single column vector for each patient (*u*_*scz*,*i*_; *i* = 1 … *N* patients), and for each drug effect (*v*_*LZP*_ and *v_MEM_*). The vectors *u* and *v* had the same length of 6 (parameters) times 180 (parcels). To assess the spatial similarity between group average maps of drug effect and individual maps of psychosis signature (map for each patient versus average map for HC group), we computed the correlation coefficient between each patient map and each drug effect map (lorazepam or memantine) as *R*_*LZP*,*i*_ = *corr*(*u*_*scz*,*i*_, *v*_*LZP*_); *i* = 1 … *N* (same for memantine).

### Statistical analyses

We computed the correlations between cortical patterns maps (e.g., psychosis signature or drug effect on dynamics parameters, receptor densities, cortical hierarchy) using Spearman’s rank correlation. To estimate the statistical significance of correlations across parcels we used spatial autocorrelation-preserving permutation tests. We generated null models using the spin rotation method (95, 96) where the cortical surface is projected into a sphere and then randomly rotated, generating permuted cortical surfaces with preserved spatial autocorrelations. We used the spherical projection of the FreeSurfer fsaverage surface for the atlas (https://github.com/rudyvdbrink/Surface_projection) to assign each parcel with the coordinates of the vertex that is closest to the center of mass of the parcel. We then randomly rotated the coordinates, and reassigned each original parcel with the value of the closest rotated parcels. We repeated this procedure 10000 times. For parcels of which the medial wall was the closest, we assigned the value of the next closest parcel. Note that we implemented the sphere rotation on the parcel-level surface, to avoid repeated values within a single permutation which often occur when re-parceling vertex-resolution surfaces (97). The same procedure was applied to assess the statistical significance of CCA (Figure 4) and of each correlation coefficient from Figure 5. We applied the same rotation angle for all parameters within a single permutation (but randomly across permutations) before vertically or horizontally concatenating them.

## Acknowledgments

We thank Karin Reimann for help with subjects’ recruitment, Annika P. Wermuth, Marlene Petersson, Barbora Schwarzova and Christiane Reissmann for help with data collection, and Anke Braun for help with experiment design.

## Author contributions

Conceptualization: AA, PJU, THD; methodology, software, investigation: AA, AT, THD; data collection: AA, AT, TG-J; data curation: AA, AT, TG-J; formal analysis, visualization: AA; writing (first draft): AA, THD; writing (review and editing): AA, AT, TG-J, PJU, THD; supervision: THD; funding acquisition: PJU, THD; project management: THD.

## Data and materials availability

Preprocessed MEG data from the pharmacological dataset will be made available upon acceptance. Preprocessed MEG data from the clinical dataset will be made available upon request. All code to reproduce the reported results and figures from the shared data will be made available on GitHub (https://github.com/DonnerLab) upon acceptance. All data are available in the main text or the supplementary materials.

## Supplementary Materials

**Figure S1:**
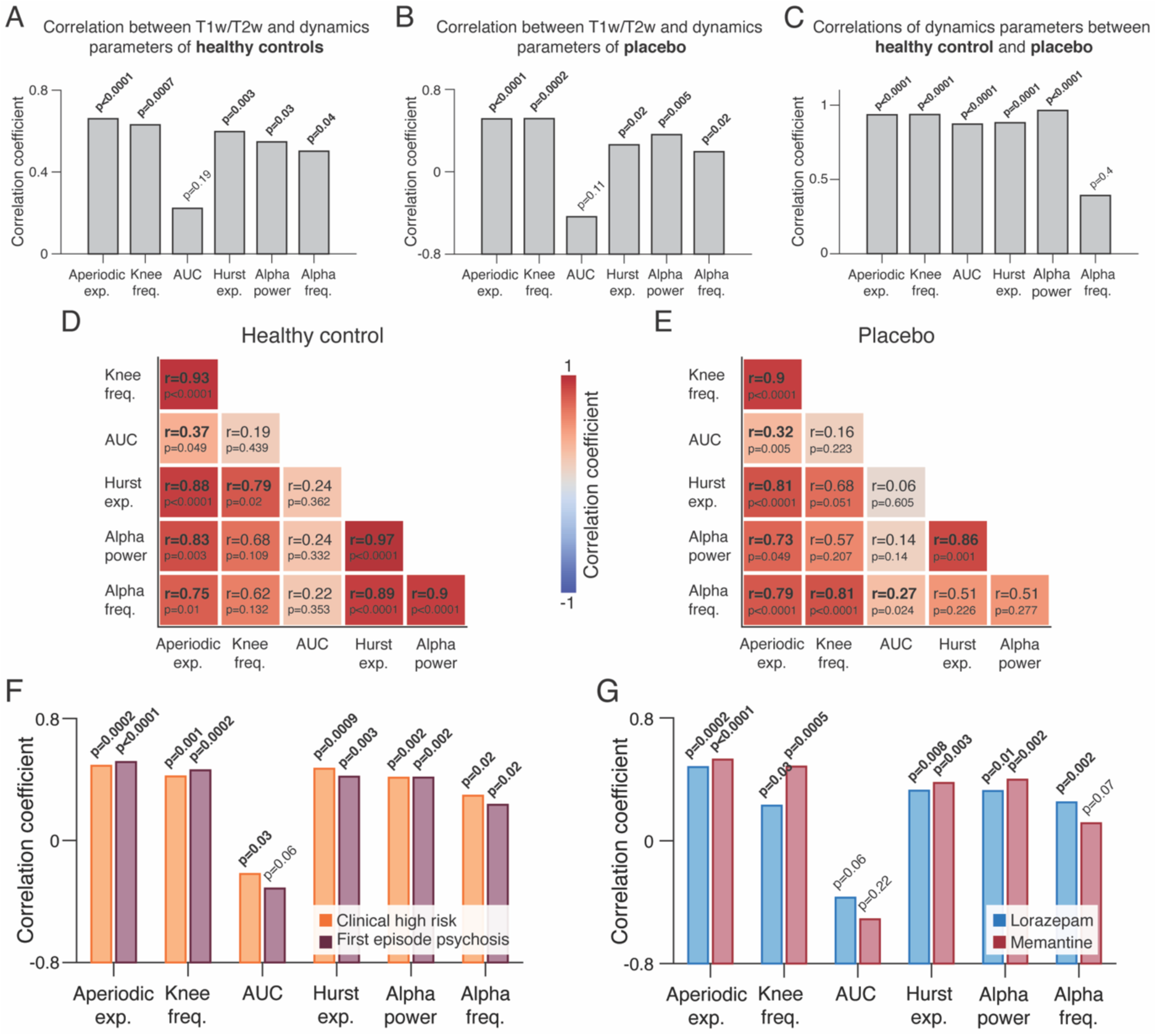
Quantification of the spatial correlation structure of spontaneous cortical dynamics. (A, **B)** Anatomical hierarchy correlates with several dynamics parameters. Across-area correlations between dynamics parameters and T1w/T2w maps. **(C)** Across parcels correlations between dynamics parameters as estimated in the placebo condition and in the healthy control group. **(D, E)** Matrices of the across-area correlations between all pairs of parameters within each sample. **(F, G)** Across-area correlations between dynamics parameters and T1w/T2w maps for clinical data (F, both clinical groups) and pharmacological data (G, both drugs). In all panels, the statistical significance of all Spearman correlation coefficients was tested using spatial autocorrelation-preserving permutation tests.

**Figure S2:**
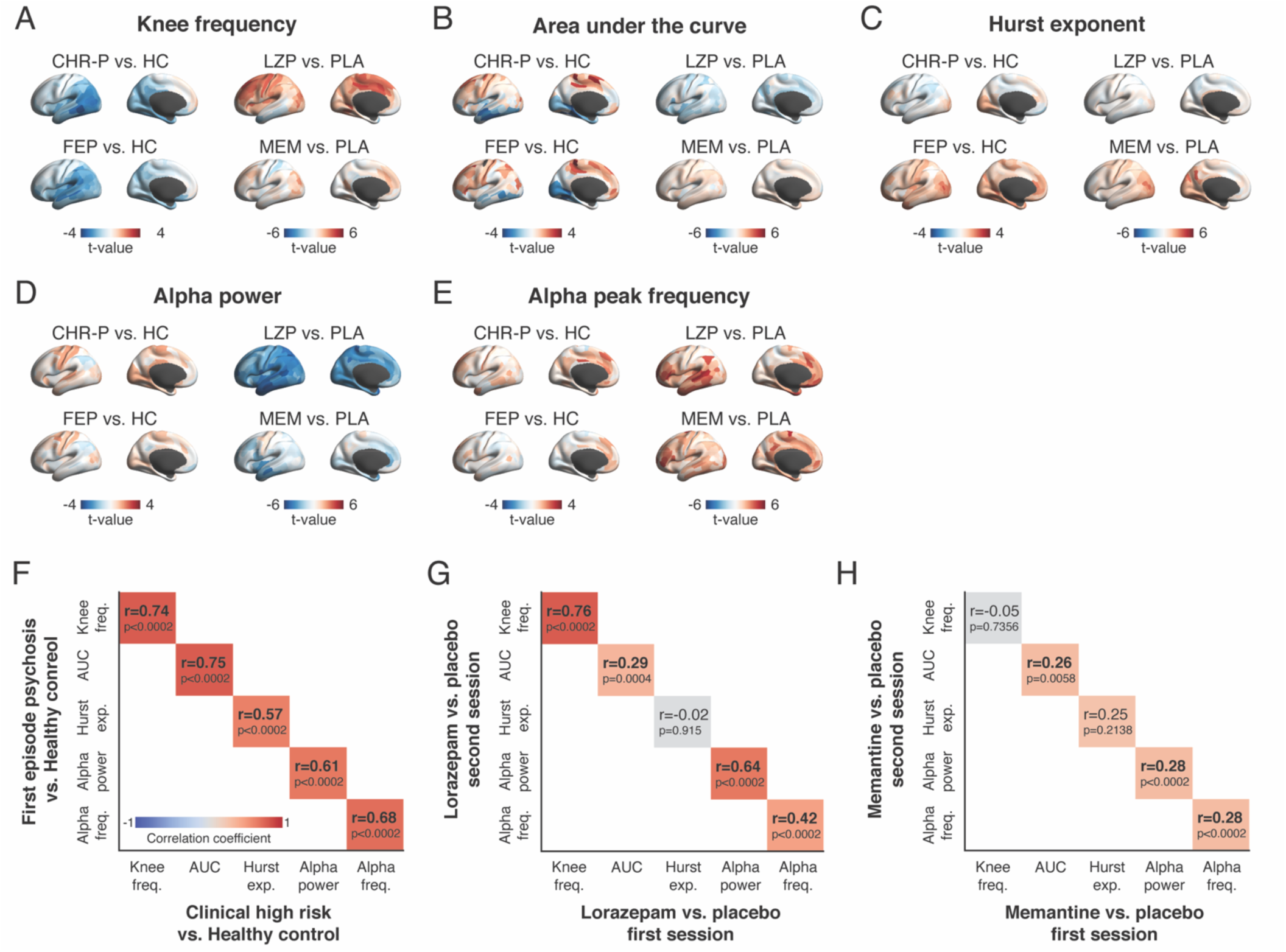
Heterogenous and reliable maps of psychosis signatures and drug effects in cortical dynamics. (A-E) Maps of drug effects or psychosis signatures on five dynamics parameters (LZP: lorazepam; MEM: memantine; CHR-P: clinical high risk; FEP: first episode psychosis; HC: healthy control). Results are presented as group level t-values. **(A)** Knee frequency, **(B)** AUC, **(C)** Hurst exponent, **(D)** Alpha power and **(E)** Alpha peak frequency. **(F)** Spatial correlations between the psychosis signature in the clinical high risk and first episode psychosis groups. **(G, H)** Spatial correlations between drug effects in the first and second repeat of the pharmacological condition, separated by 1-5 weeks. (G) Lorazepam, (F) memantine. All correlations are Spearman’s correlation coefficients and all p-values were estimated using spatial autocorrelation-preserving permutation tests.

**Figure S3:**
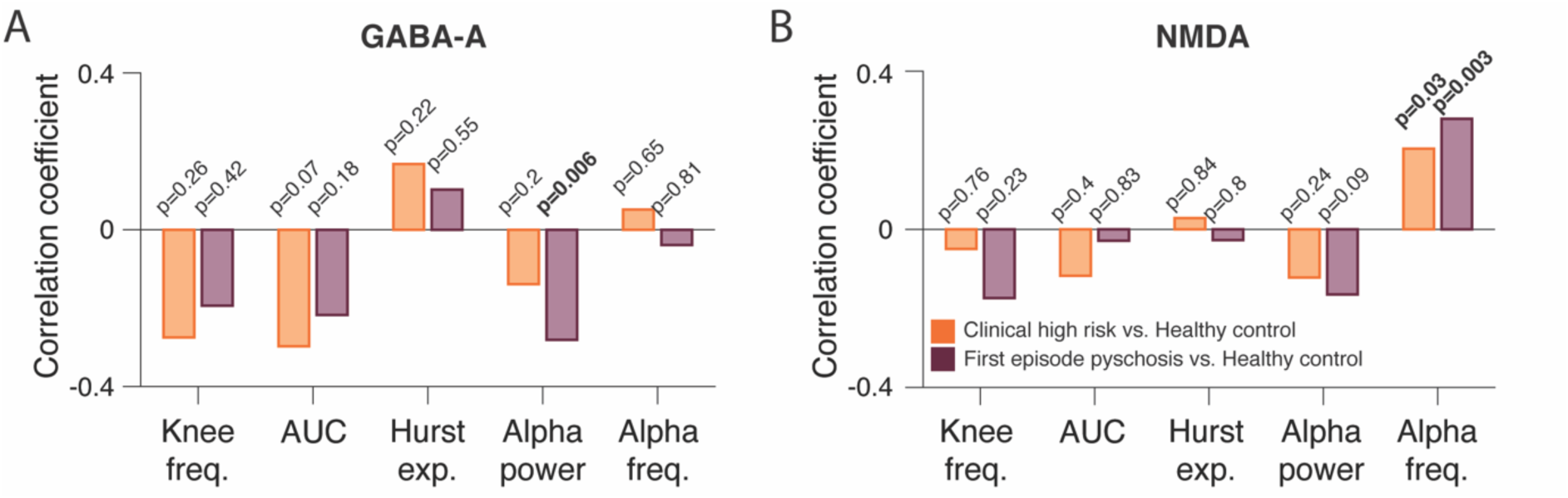
Similarity between maps of psychosis signatures on dynamics parameters and maps of GABA-A or NMDA receptor density levels. (A, **B)** Spatial correlations between the psychosis signature on dynamics parameters and GABA-A (A) or NMDA (B) receptor densities. All correlations are Spearman’s correlation coefficients and all p-values were estimated using spatial autocorrelation-preserving permutation tests.

**Figure S4:**
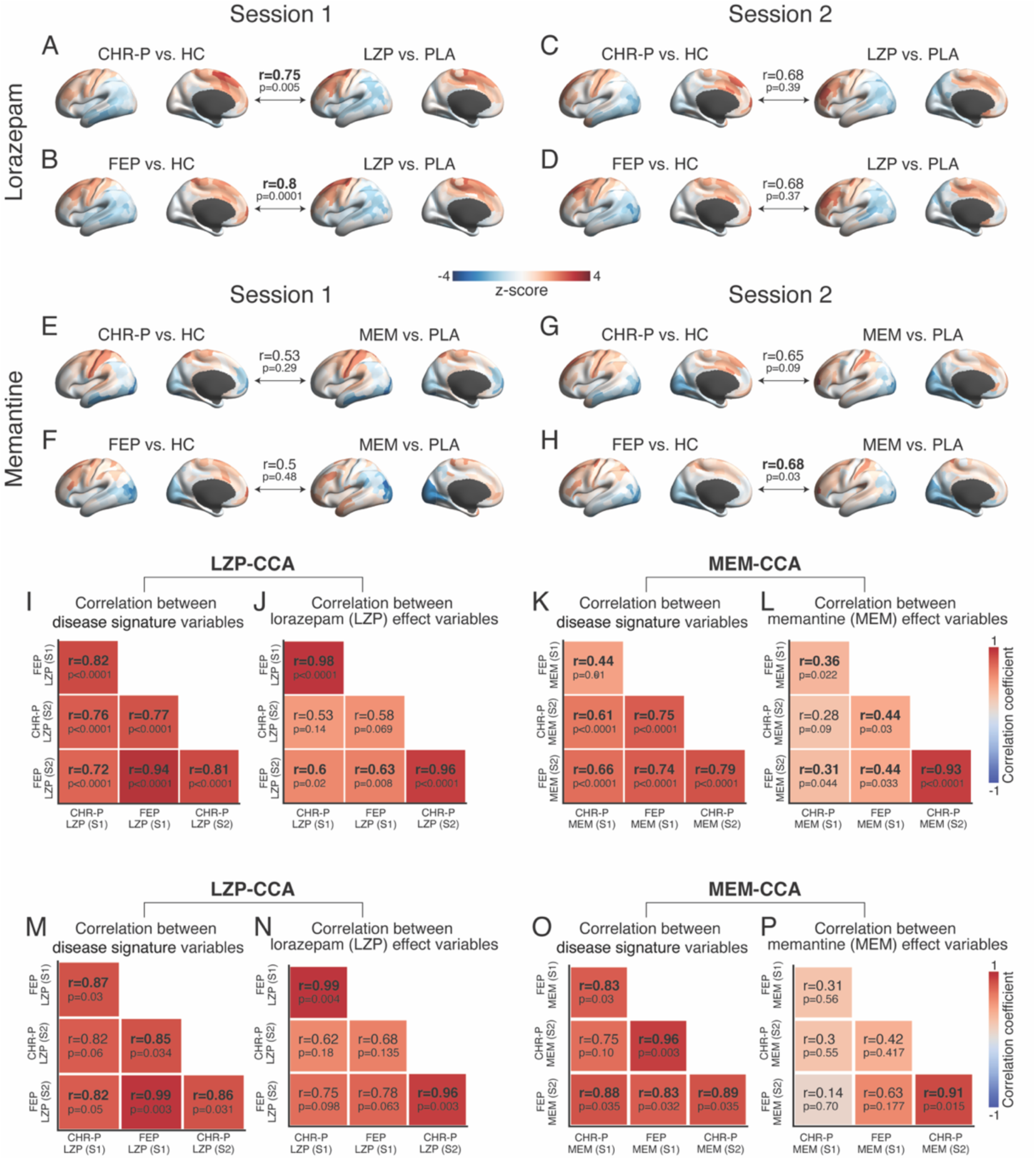
CCA results were highly reproducible across repeats of pharmacological conditions and for different clinical groups. (A-H) spatial patterns of the first canonical variable of the psychosis signature (left) or drug effect (right) and the corresponding canonical correlation between them (horizontal arrows). CCA was repeat for each pair of drug experimental sessions: (session 1, session 2) and clinical group (CHR-P vs. HC, FEP vs. HC; CHR-P: clinical high risk; FEP: first episode psychosis; HC: healthy control). **(I-J)** Correlation matrices between all pairs of canonical variables in (A-H), separated by drug (I, J: lorazepam; K, L: memantine), and canonical variable (I, K: psychosis signature; J, L: drug effect). All correlations are Spearman’s correlation and p-values were estimated using spatial autocorrelation-preserving permutation tests. **(M-P)** same as (I-L) but for the CCA loading patterns.

**Figure S5:**
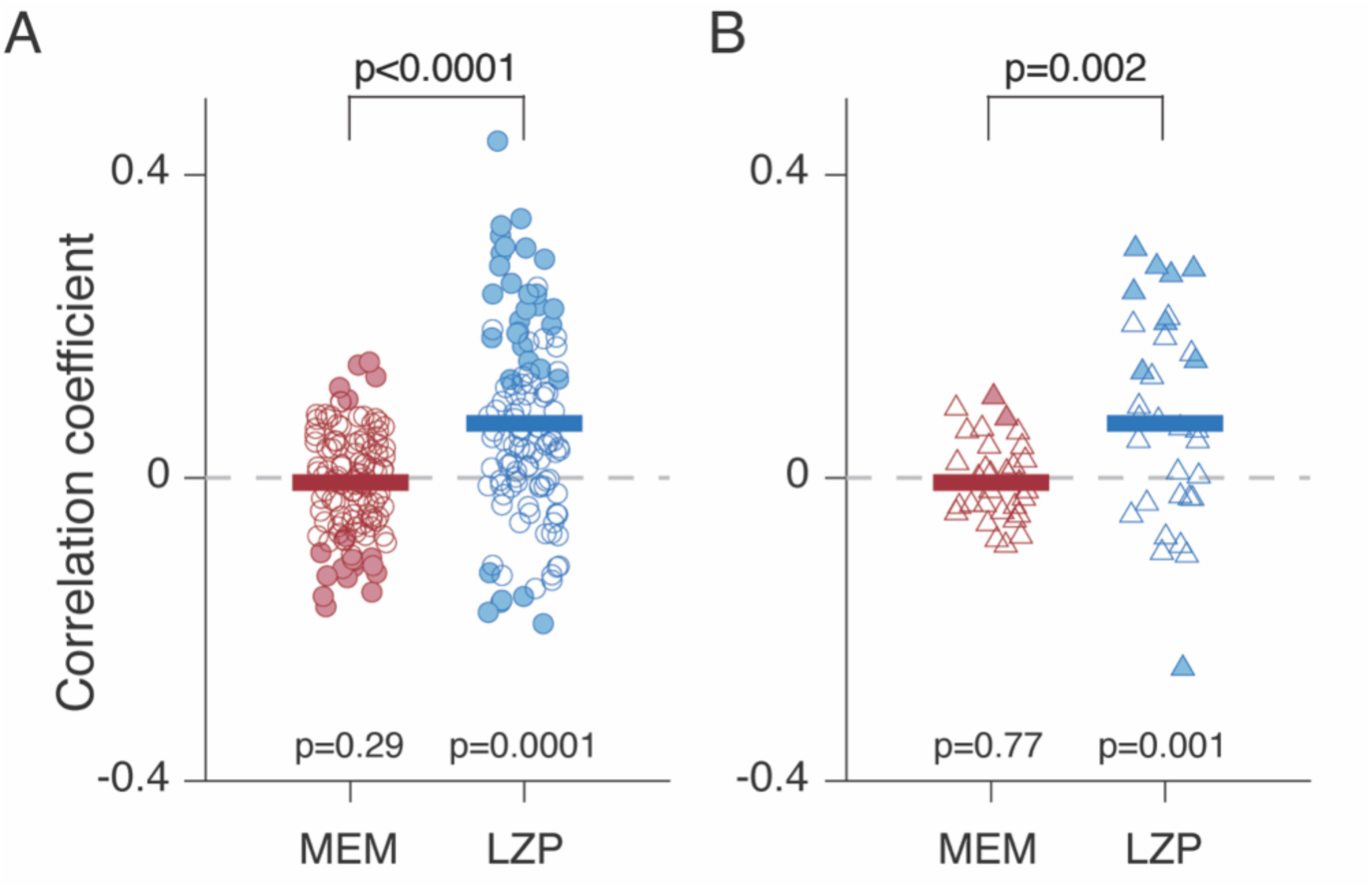
Spatial similarity between group-average drug effects and individual psychosis signatures (individual patient vs. HC average). Each dot represents the level of pattern similarity for a single patient and a given drug effect (LZP or MEM). Filled symbols, individual patients with statistically significant pattern similarity with a given drug effect. Same data as Figure 5B but split by CHR (A) and FEP (B) groups, with separate tests for both. The results are identical to the pooled analysis results in Figure 5B.

**Figure S6:**
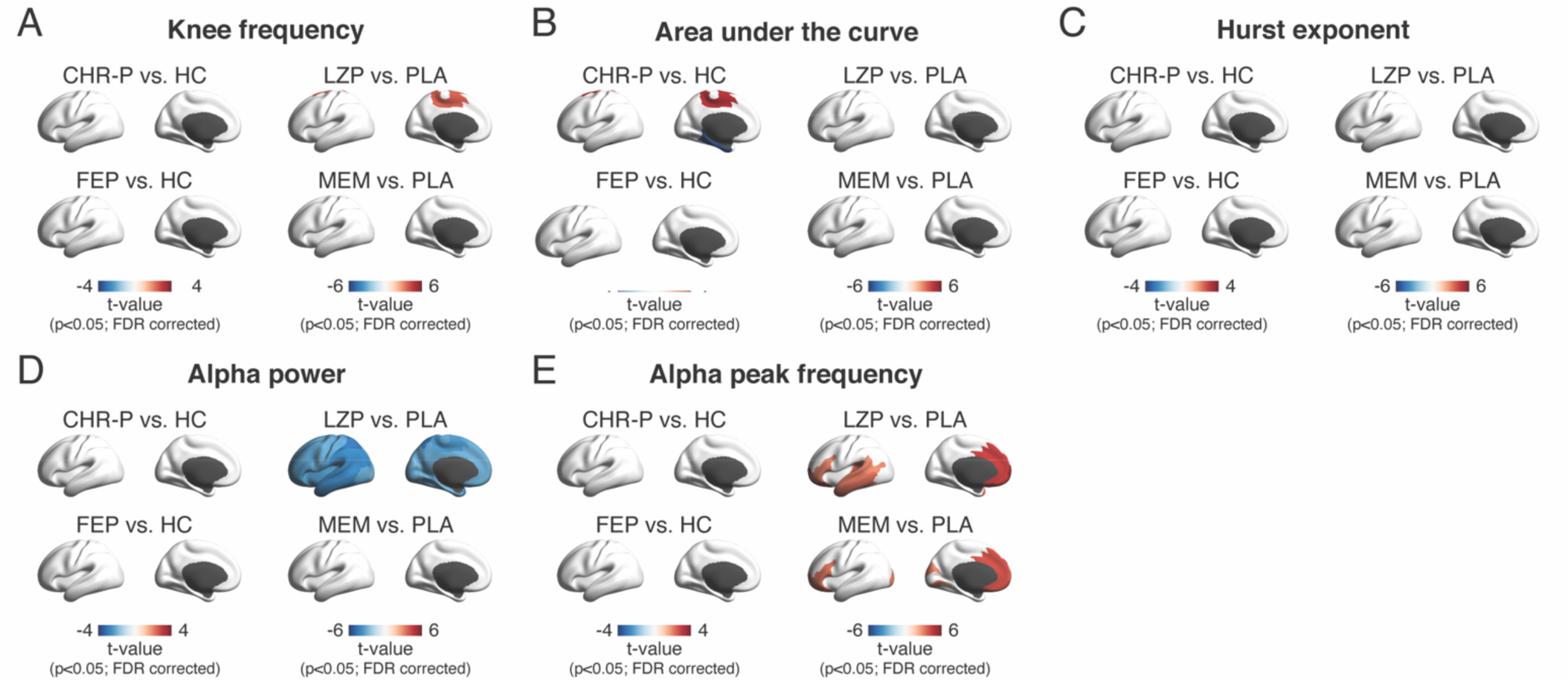
Weak psychosis signatures or drug effects in local cortical dynamics. (A-E) Maps of psychosis signatures (CHR-P vs. HC or FEP vs. HC; left) or drug effects (lorazepam vs. placebo or memantine vs. placebo; right) on **(A)** knee frequency, **(B)** AUC, **(C)** Hurst exponent, **(D)** alpha power, and **(E)** alpha peak frequency. The widespread reduction in alpha power under LZP in panel D is a well-documented phenomenon ^98-100^. CHR-P: clinical high risk; FEP: first episode psychosis; HC: healthy controls; LZP: lorazepam; MEM: memantine. All maps are presented as group level t-value of significant parcels only (p<0.05, FDR correction). All p-values were assessed with two-sided permutation tests.

